# Mapping and rewiring the *MYBPC3* promoter for rescue of haploinsufficiency driven hypertrophic cardiomyopathy

**DOI:** 10.64898/2026.09.20.752987

**Authors:** Aaron Renberg, Savannah Coppersmith, Owen Merritt, Yao-Chang Tsan, Esraa Ismail, Jennasea Licata, Maya Sheth, Revant Gupta, Anshul Kundaje, Sabrina Friedline, Hayden C. Helms, Elizabeth D. Hughes, Zachary T. Freeman, George Powell, James S. Ware, Joshua Meisner, Eric M. Smith, Adelaide Tovar, Jesse M. Engreitz, Jacob Kitzman, Adam S. Helms

**Affiliations:** Department of Internal Medicine, Division of Cardiovascular Medicine, University of Michigan, Ann Arbor, Michigan, USA; Cellular and Molecular Biology Program, University of Michigan Medical School, Ann Arbor, Michigan, USA; Department of Molecular and Integrative Physiology, University of Michigan, Ann Arbor, Michigan, USA; Department of Genetics, Stanford University School of Medicine, Stanford, CA, USA; Basic Sciences and Engineering Initiative, Betty Irene Moore Children’s Heart Center, Lucile Packard Children’s Hospital, Stanford, CA, USA; Department of Bioengineering, Stanford University, Stanford, CA, USA; Department of Computer Science, Stanford University, Stanford, CA, USA; Transgenic Animal Model Core, University of Michigan, Ann Arbor, Michigan, USA; National Heart and Lung Institute and MRC Laboratory of Medical Sciences, Imperial College, London, UK; Department of Pediatrics, Division of Pediatric Cardiology, University of Michigan, Ann Arbor, Michigan, USA; Department of Human Genetics, University of Michigan, Ann Arbor, Michigan, USA; Gene Regulation Observatory, Broad Institute of MIT and Harvard, Cambridge, MA, USA; Novo Nordisk Foundation Center for Genomic Mechanisms of Disease, Broad Institute of MIT and Harvard, Cambridge, MA, USA; Stanford Cardiovascular Institute, Stanford University, Stanford, CA, USA

**Author notes:** denotes corresponding author Corresponding Author: Adam S. Helms, 1500 E. Medical Center Drive Ann Arbor, MI 48019.

## Abstract

Autosomal dominant loss-of-function variants in the gene *MYBPC3* are, collectively, the most common genetic cause of hypertrophic cardiomyopathy (HCM) and are a prototype of haploinsufficient human disease. Typical for haploinsufficiency-associated genes, hundreds of unique loss-of-function pathogenic variants have been reported for *MYBPC3* – therapeutic gene editing to correct each of these variants poses major regulatory and logistical hurdles. Upregulating wild-type allele expression could offer a generalizable therapeutic strategy, but the capacity to modulate native *MYBPC3* transcription is unknown. Here, we present a variant-agnostic approach to rescue haploinsufficiency by mapping and rationally redesigning the *MYBPC3* promoter. Using massively parallel reporter assays (MPRAs) in human induced pluripotent stem cell-derived cardiomyocytes, we performed saturation mutagenesis of the *MYBPC3* promoter at single base-pair resolution. This defined a new class of clinically relevant noncoding loss-of-function variants while revealing essential cis-regulatory grammar anchored by key transcription factor binding sites (TFBSs). Furthermore, by systematically screening thousands of variant combinations, modular promoter elements, and heterologous TFBS insertions, we identified synergistic sequence edits that drive robust increases in *MYBPC3* expression. Together, our findings improve the clinical interpretation of noncoding variants and establish a scalable blueprint for promoter editing to treat *MYBPC3*-associated HCM and other haploinsufficient diseases.

## Introduction

Autosomal dominant loss of function variants in the gene *MYBPC3* are, collectively, the most common genetic cause of hypertrophic cardiomyopathy (HCM) and are a prototype of haploinsufficient human disease.^1,2^ *MYBPC3* encodes cardiac myosin binding protein C3 (cMyBP-C), which regulates cardiac sarcomere function, and *MYBPC3* loss of function results in dysregulation of actin-myosin sliding filament interactions.^3,4^ The lifetime disease burden is high – many patients develop heart failure and arrhythmias.^5^ A small proportion develop advanced heart failure necessitating heart transplant.

There are currently no FDA-approved treatments for *MYBPC3*-HCM. While outflow obstruction, a consequence of HCM, can be mitigated by a new class of myosin inhibitor medications, treatments for the fundamental “nonobstructive” myopathy component of HCM have been limited.^6–9^ Gene replacement therapy delivered by adeno-associated virus (AAV) has been advanced as a strategy for *MYBPC3*-HCM. However, *MYBPC3* is a very highly expressed gene, posing a challenge for maintenance of the high levels of protein required to maintain sarcomere stoichiometry. Additionally, the durability of expression in humans with a single dose of AAV based gene therapy is unknown, and current AAV platforms do not allow re-dosing due to antibody neutralization. Individual variant correction through emerging technologies, such as base or prime editing, could offer a more durable curative treatment, but the time, expense, and regulatory hurdles to confront hundreds of unique *MYBPC3* truncating variants is considerable.^10^ Notably, despite *MYBPC3* variants being established as a common genetic cause of HCM ∼3 decades ago, the transcriptional regulation of *MYBPC3* has been scarcely studied.^11^

Here, we developed a variant-agnostic and durable approach to rescue haploinsufficiency due to *MYBPC3* loss of function. Since *MYBPC3* truncating variants are etiologically similar and do not exhibit dominant negative effects, we reasoned that upregulating *MYBPC3* transcription could provide a universal approach to restore normal levels of *MYBPC3* mRNA. Using massively parallel reporter assays (MPRAs) conducted in cardiomyocytes differentiated from induced pluripotent stem cells (iPSC-CMs), we first mapped the *MYBPC3* promoter at single base pair resolution with saturation mutagenesis. These MPRAs revealed loss of function variants clustering in transcription factor binding sites (TFBSs), enabling us to decipher indispensable promoter elements while also defining a new class of noncoding loss of function variants relevant for clinical genetic diagnosis in HCM. Additionally, we identified gain of function variants and performed iterative combinatorial variant effect testing to identify variant combinations with additive effects. We further systematically assessed integration of other promoter and TFBS elements to test effects on expression in the *MYBPC3* promoter context. This work demonstrates that promoter editing is capable of rescuing haploinsufficiency as a variant agnostic approach, even for a highly expressed gene such as *MYBPC3*.

## Results

### Mapping the *MYBPC3* promoter with saturation mutagenesis reveals loss of function variants clustered in transcription factor binding sites

The *MYBPC3* promoter coordinates were defined based on ENCODE^12^ annotations in the UCSC genome browser^13^. We selected a 456 bp region extending from -428 bp from the transcriptional start site (TSS) to +28 bp into the 5’ UTR for mapping. This region encompasses the core *MYBPC3* promoter that we previously demonstrated is necessary for *MYBPC3* transcription.^1^ To map the *MYBPC3* promoter, while simultaneously determining single nucleotide variant (SNV) effects, we performed synthetic saturation mutagenesis using oligo libraries and lentiviral MPRAs (**Figure 1A**).^14^ Lentiviral MPRA libraries were transduced into iPSC-CMs at a scale to obtain at least 10 barcodes per variant per replicate. 93% of all possible SNVs were measured. An average of 84% of variants had at least 25 unique barcodes captured (**Supplemental Figure S1A)**. Concordance among replicates and between cell lines was high (mean pairwise *r* = 0.71, 95%CI = [0.58,0.85] for library 1 replicates; mean pairwise *r* = 0.80, 95%CI = [0.71,0.90] for library 2 replicates, *r* denotes Pearson correlation, **Supplemental Figure S1B-C**). A compendium of all variants and effect sizes is shown in **Supplemental Table S1**.

**Figure 1:**
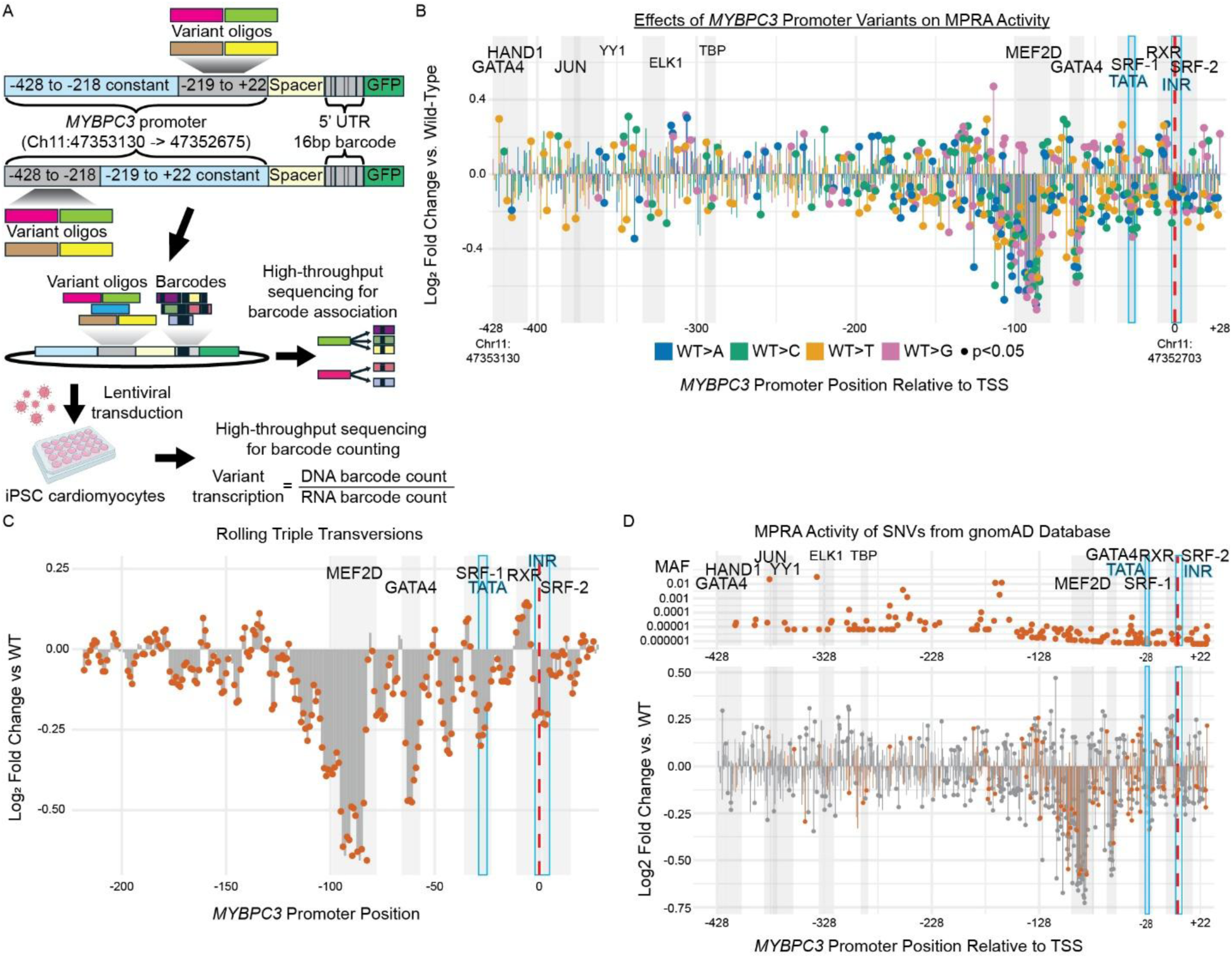
Saturation mutagenesis of *MYBPC3*’s promoter in an MPRA. MPRA was performed using a variant library containing each possible SNV in the *MYBPC3* 456bp core promoter sequence. **(A)** MPRA construct design consisted of pools of SNV, triple transversions, or wild type (WT) sequence oligos placed next to remaining *MYBPC3* promoter sequence as a constant sequence. A 20bp constant spacer sequence followed, then a random 16bp barcode in the 5’ UTR of *GFP*. Our lentiMPRA workflow was adapted from Gordon et al.^14^ Each variant oligo was randomly paired with a unique 16bp barcode in a modified pLS-SceI backbone. High throughput sequencing of the plasmid pool was used to build a dictionary of variant-barcode pairs. Induced pluripotent stem cell-derived cardiomyocytes were transduced with lentivirus packaged from this plasmid pool. DNA and RNA were extracted and then high throughput sequencing targeting the barcode was used to obtain barcode counts which were interpreted using the variant-barcode dictionary. Normalizing the barcode counts in the RNA (expression) to the counts in the DNA (viral insertion events) provides a measure of variant effect on transcription. See Methods for further details. **(B)** RNA/DNA barcode count ratio was calculated and compared to wild-type. Many variants were found to have significant effects on transcription (p-adj <0.05, Wilcoxon test with Benjamini-Hochberg correction). **(C)** MPRA activity of rolling triple transversions spanning the 3’ half of the MYBPC3 promoter. For each position, mean activity for all variants spanning that position was compared to wild-type activity. **(D)** Mean allele frequencies in gnomAD for MYBPC3 promoter variants above their activity in the MPRA. 69 variants with significant effects in the MPRA appear in gnomAD.

Of the 1276 variants included in the analysis, 421 variants (33.0%) significantly impacted transcription (p-adj <0.05, **Figure 1B, Supplemental Table S1**). Of these, 146 (34.7%) increased transcription with a median log_2_ fold-change compared to wild-type (LFC_WT_) of 0.142 (+10.3%, IQR=[0.109,0.189]) and 275 (65.3%) decreased transcription with a median LFC_WT_ of -0.185 (-12.0%, IQR = [-0.296,-0.125]). Variants that reduced transcription were significantly enriched in TFBSs including MEF2 and GATA4 (OR = 1.96, 95%CI = [1.47,2.62], p<0.0001). Variants with positive effects were not significantly enriched in TFBSs (OR = 0.73, 95%CI = [0.47,1.09], p=0.126). The mean effect size weakly correlated with vertebrate conservation by position (ρ = 0.183, p < 0.0001, Spearman correlation, **Supplemental Figure S2A**). Of the variants with significant effects, 69 (16%) are reported as present in gnomAD (**Figure 1D**). Effect size of these variants ranged from -33% to +19% and are reported, along with their gnomAD allele frequencies, in **Supplemental Table S1**. Most variants with significant effects present in gnomAD were rare (62 of 69 with MAF<1e-5).

As an additional validation, we also mapped the core promoter with rolling 3 bp transversions spanning from -217 to +28 relative to the TSS. Variant effects with this approach highly correlated with SNV effects at each position (*r* =0.82, p<2.2e-16, **Supplemental Figure S2B, Supplemental Table S2**). This approach similarly mapped TFBSs with high signal:noise and notably required fewer oligonucleotides in the pooled libraries (**Figure 1C**).

Our libraries were designed with an additional 5’ enhancer to increase overall signal to overcome a limitation observed in a similar study.^15^ We also tested a portion of the SNVs without this enhancer. While the enhancer did increase transcription (1.08-fold mean increase, p<0.0001, Wilcoxon test), its presence did not reduce MPRA measurement variance (CV=0.58 with vs 0.57 without, p<0.01, **Supplemental Figure S2C-D**).

### Variants with significant effects act through disruption or creation of key cardiac TFBSs

To better understand the mechanisms by which the variants tested in our MPRA influence transcription, we analyzed their effects on empirically identified TFBSs. We used the R package motifmatchr^16^ to scan a 31 bp window centered on each SNV for TFBS motifs and calculated the predicted impact of that SNV on binding affinity (delta score, see Methods and **Supplemental Figure S3A**). We annotated the *MYBPC3* promoter sequence with these TFBSs based on the JASPAR^17^ track in the UCSC Genome Browser^13^. Of these TFBSs, GATA4, and MEF2D variants decreased transcription compared to non-TFBS variants (**Figure 2A**). Variants in GATA4, MEF2D, and the TATA box were individually more likely to have a significant effect on transcription than non-TFBS variants, highlighting their vulnerability to SNVs (**Figure 2B**). Additionally, simultaneously replacing GATA4, MEF2D, and the TATA box with randomized sequences resulted in a 37.5% reduction of transcription compared to wild-type (95%CI = [36.1%,38.9%]), further supporting these sequences’ essential role (**Supplemental Table S3**). Correlating delta score with our MPRA data can empirically identify TFs acting as activators or repressors. Correlations for individual TFs confirm that GATA4, MEF2D, the first SRF motif, and the TATA box act as activators and that RXRA acts as a repressor (**Figure 2C, Supplemental Figure S3B**). Motifmatchr analysis of highly positive SNVs that were not within an annotated TFBS reveals that the two most positive SNVs both create *de-novo* cardiac-relevant TFBSs – - 70G>C (LFC_WT_ = 0.29) creates a weak GATA4 motif (delta score = 0.06), and -114C>G (LFC_WT_ = 0.47) creates a new JUND/FOSL2 motif (delta score = 8.99).

**Figure 2:**
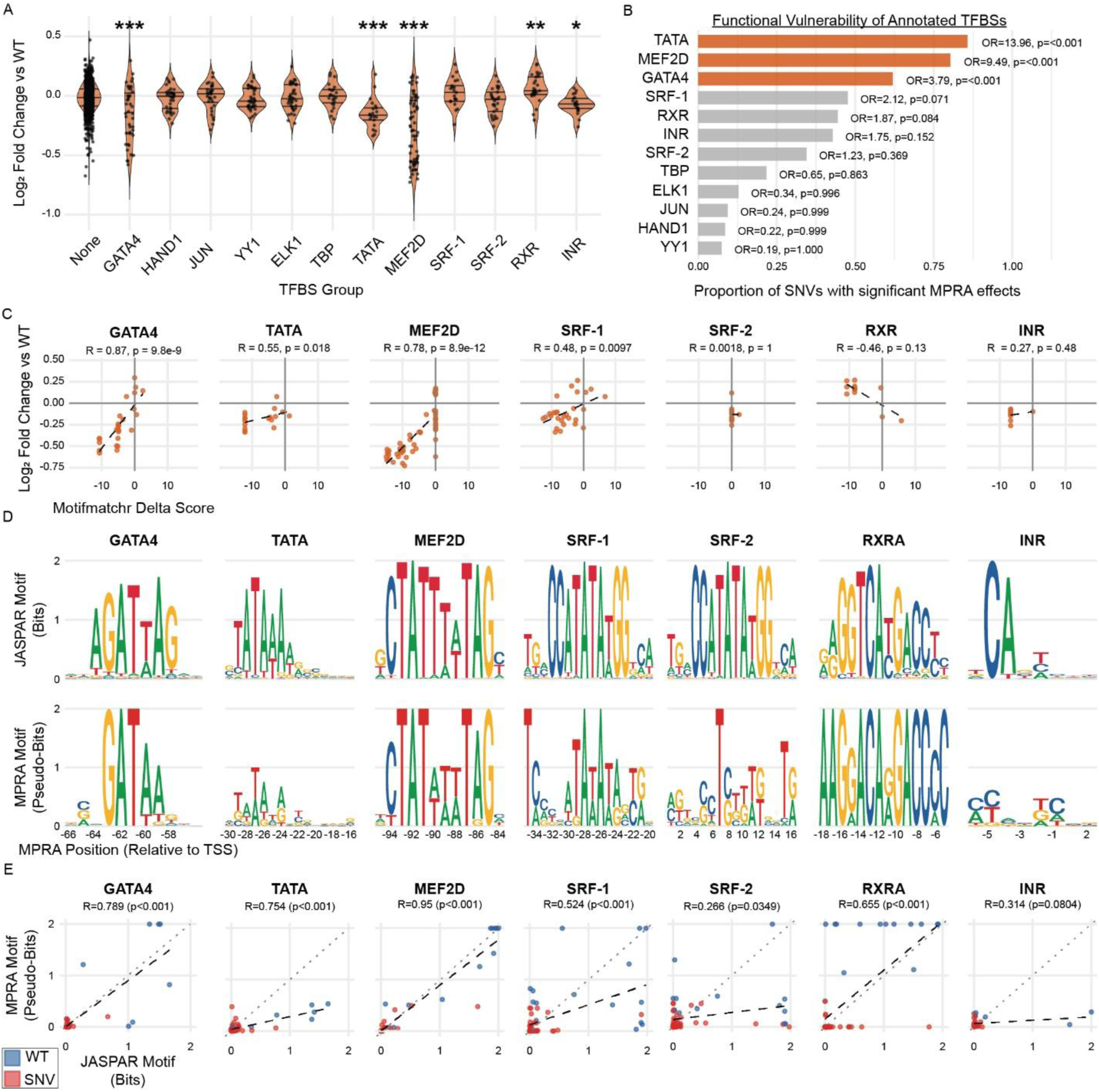
Transcription factor binding site motif analysis demonstrates that key cardiac transcription factors are vulnerable to SNVs. **(A)** MPRA data for SNVs within each annotated TFBS. Variants within GATA4, and MEF2D decrease expression compared to non-TFBS SNVs, consistent with their roles as cardiac-active transcription factors. Variants within the TATA box, and initiator sequence (INR) similarly reduce expression compared to the non-TFBS SNVs. Variants within RXR increase expression compared to non-TFBS SNVs, suggesting it acts as a repressor in this context. (Dunnett test, *p<0.05, **p<0.01, ***p<0.001) **(B)** Vulnerability analysis for each annotated TFBS. Variants within GATA4, MEF2D, and the TATA box are enriched for significance (Fisher’s exact test, p<0.001). **(C)** Predicted SNV impact on individual TFBS binding affinity (Delta Score) was calculated using motifmatchr (see Methods) and compared to impact on transcription in the MPRA (Pearson correlation). Positive correlation indicates that TF is acting as an activator while negative correlation suggests it is acting as a repressor. See **Supplemental Figure S3B** for additional TFBSs. **(D)** Logo-style motifs for TFBSs were generated empirically from MPRA SNV data by calculating a contribution score for each base at each position (see Methods) and are shown beneath their canonical motifs from JASPAR^17^. See **Supplemental Figure S3C** for additional TFBSs. **(E)** The contribution score for each base calculated in generating the logo-style motifs was compared to the contribution scores from the canonical JASPAR motif (Pearson correlation). MEF2D, GATA4, the TATA box, and the first SRF site empirically generated motifs are highly correlated with their canonical motifs. See **Supplemental Figure S3D** for additional TFBSs.

We empirically generated logo-style binding motifs, which reflect the local sequence preferences at putative TFBSs by calculating a contribution score for each base using the MPRA (**Figure 2D**). MPRA-derived motifs were highly concordant with canonical motifs for critical TFBSs (**Figure 2E**). The MPRA-generated motif for SRF-1 showed the TATA motif as the most critical, with little preference for the flanking CC and GG expected for the putative SRF binding motif, suggesting that the sensitivity of this region to SNVs is likely more due to its role as the TATA site than a true SRF binding site. Thus, the MPRA-based motif analysis appears to detect context-dependent behavior beyond sequence-only based predictions. When we performed these analyses for other JASPAR annotated TFBSs (e.g., JUN, YY1), there was a lack of evidence for these as important TFBS sites for the *MYBPC3* promoter (**Supplemental Figures S3C-D**).

Additionally, promoter mapping with 3 bp transversion disruptions identified two regions (-177 to -163 and -157 to -143) for which disruptions affected expression (**Figure 1C**). Analysis with motifmatchr identified an overlapping pair of NFIX sites (-160 to -152 and -156 to - 148, delta score = -10.92) that have large decreases in delta score when disrupted by transversions. Corroborating these data, motifmatchr analysis of SNV data identifies -155C>T as disrupting an NFIX site (delta score = -8.8).

Taken together, these analyses decipher the key regulatory grammar that drives transcription of the *MYBPC3* promoter.

### *In silico* saturation mutagenesis of the *MYBPC3* promoter using ChromBPNet and AlphaGenome models reflects MPRA data for most impactful SNVs

We next assessed how well our MPRA data correlated with machine learning (ML) model predictions. We compared our MPRA data to predictions from both ChromBPNet and AlphaGenome ML models. From ChromBPNet, 25 deep-learning models trained on DNase (N=10) and ATAC-seq (N=15) data from human left ventricle samples were used to derive predictions for *in silico* saturation mutagenesis of the *MYBPC3* promoter.^18,19^ The ChromBPNet models’ predictions were highly correlated to each other (*r =* 0.763, 95%CI = [0.760,0.766], p < 0.0001, **Supplemental Figure S4A**). The read count predictions were used to calculate a predicted LFC_WT_ which was compared to MPRA data (**Figure 3A**). The predicted LFC_WT_ correlated overall with the MPRA data (*r* = 0.64, 95%CI = [0.60,0.67], p< 0.0001, **Figure 3B**). The model had higher correlations within the high impact TFBSs, as follows: MEF2D (*r* = 0.85, 95%CI = [0.78,0.91], p < 0.0001), GATA4 (*r* = 0.66, 95%CI = [0.44,0.80], p < 0.0001), SRF-1 (*r* = 0.55, 95%CI = [0.29,0.73], p = 0.0002), and RXR (*r* =0.61, 95%CI = [0.31,0.81], p=0.0006). The remaining TFBSs were not statistically significantly correlated. The model predicted lower impact of variants farther away from the transcriptional start site even though the MPRA data shows some of these variants had significant effects on transcription. There was no statistical difference in median total absolute residual between ATAC-seq trained models and DNAse-trained models (0.078 vs 0.077, p=0.24, Wilcoxon test). Notably, the models tended to underpredict absolute effect of variants and predicted the opposite effect than observed for 25.4% of the variants with statistically significant MPRA effects (12.8% MPRA+ ChromBPNet-, 12.6% MPRA-ChromBPNet+). A possible explanation of this is bias in the model’s training data toward peaks centered on the TSS.^18^ Predictions from AlphaGenome, another model derived using data outputs of ATAC-seq, CAGE, and DNAse assays, performed similarly. Predictions from AlphaGenome moderately correlated with MPRA data (*r* = 0.60, p<0.0001, **Supplemental Figure S5A**) and strongly correlated with predictions from ChromBPNet (*r* = 0.85, p<0.0001, **Supplemental Figure S5B**).

**Figure 3:**
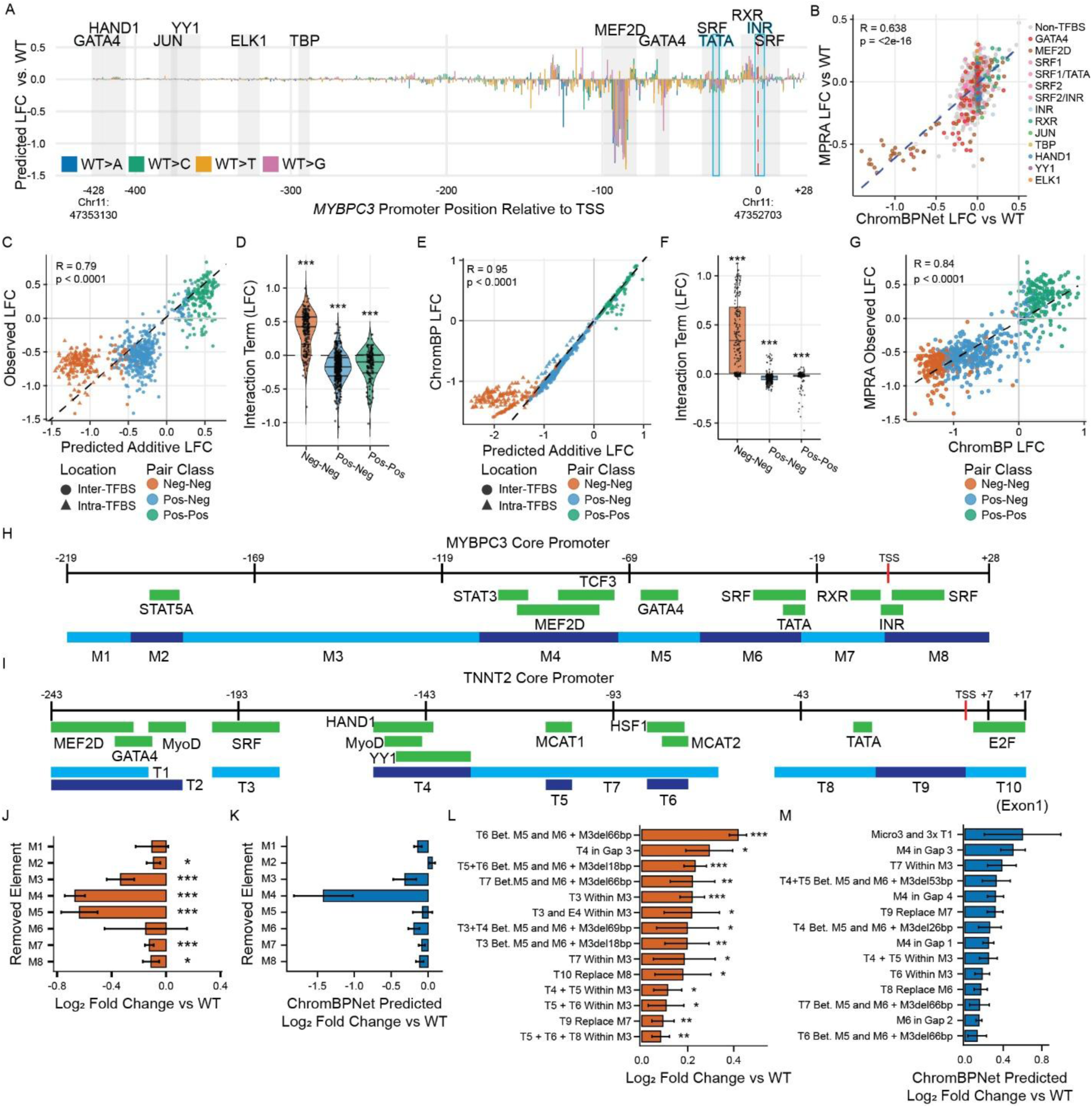
Linear modeling analysis of paired SNVs to determine additive or epistatic Effects. **(A)** 25 ChromBPNet models were trained on different DNase and ATAC-seq datasets from human left ventricle samples. These models were used to perform in silico mutagenesis on the *MYBPC3* promoter. (LFC_WT_ for simulated ATAC-seq read counts) **(B)** These SNV effect predictions were correlated to the LFC_WT_ observed in the MPRA experiment. (Pearson correlation, p<0.0001) **(C)** The SNVs with the 20 largest positive and negative MPRA effects were selected for analysis of the effects of pairs of SNVs in an MPRA. A weighted linear model was created for each possible pair among these 40 variants using the single SNV effects to predict the paired effect assuming additive effects. These predictions were highly correlative with actual MPRA activity (R=0.79, p<0.0001, Pearson correlation). **(D)** The interaction term of the model represents any epistatic effect. All three pair classes display a sub-additive behavior. **(E)** The ChromBPNet models predicted LFC_WT_ for paired SNVs were compared to the summed predicted effect of the two individual SNVs. **(F)** The interaction term for the ChromBPNet predictions for paired SNVs were smaller in magnitude than observed in the MPRA, but followed the same pattern, with statistically significant sub-additive behavior predicted. **(G)** The observed MPRA LFC_WT_ and ChromBPNet predicted LFC_WT_ were highly correlated. (R=0.84, p<0.0001) **(H-I)** The *MYBPC3* and *TNNT2* core promoters (*MYBPC*3: chr11: 47352703-47352920, *TNNT2*: chr2:199141718-199141737) were divided into 8 and 10 elements, respectively, based on TFBS annotations and regions of high evolutionary conservation. **(J)** Effects on transcription of replacing each individual element with random sequence compared to wild-type promoter. **(K)** ChromBPNet predicted LFC_WT_ for the same promoter element replacements. **(L)** MPRA was performed using 68 rationally designed *MYBPC3* and *TNNT2* hybrid promoters. The 14 shown had statistically significant positive effects on transcription compared to the full WT *MYBPC3* promoter. (See **Supplemental Figure S6B** for all promoters tested.) **(M)** ChromBPNet predicted LFC_WT_ was generated for the same hybrid promoters. The 14 hybrid promoters shown here were predicted to be the most transcriptionally active. Significance: *FDR<0.05, **FDR<0.01, ***FDR<0.001, Wilcoxon rank sum test with Benjamini-Hochberg correction and 95%CI unless otherwise noted.

The high concordance of the ChromBPNet predictions and the MPRA for variants near the TSS suggests this approach could be used to prioritize regions of interest for study. Nevertheless, the incomplete concordance both in effect direction and magnitude highlight the need for continued empirical assessment of noncoding elements and further development of machine learning methods.

### Combinatorial effects of *MYBPC3* promoter variants

We next determined additivity for *MYBPC3* promoter SNVs. We performed an MPRA that tested all possible pairs of the variants with the 20 strongest positive or negative effects (40 total) from the promoter saturation mutagenesis mapping above (**Supplemental Figure S4B**). These variants included 17 MEF2D variants, one GATA4 variant, two SRF variants, five RXR variants and 15 variants that were not within a TFBS. Data for 742/780 possible pairs (95%) passed MPRA quality filtering (**Supplemental Table S4**). A weighted least squares linear regression was applied to the replicate-level MPRA LFC_WT_ for each pair of single SNVs to predict the effect of the combined pair; the coefficient of the interaction term indicated potential non-additive effect (i.e., epistasis). Overall, predictions from the linear regression strongly correlated with the experimental MPRA LFC_WT_ across variant combinations (*r* = 0.79, p<0.0001, **Figure 3C**). This correlation did not meaningfully change when pairs of SNVs in the same TFBS are excluded (*r* = 0.80, p<0.0001, **Supplemental Figure S4C**). We also performed regressions for SNV combinations classified by shared or opposing effect directions. The pairs with one positive- and one negative-effect SNV (Pos-Neg) were the most tightly correlated (*r* = 0.61, p<0.001), but pairs with two negative-effect SNVs (Neg-Neg) and two positive-effect SNVs were also correlated (*r* = 0.29, p = 0.005 and *r* = 0.26, p = 0.0007, respectively). Neg-Neg pairs demonstrated epistasis, with actual LFC_WT_ less negative than predicted (Median LFC_WT_ - LFC_Predicted_ = 0.425, 95%CI = [0.339,0.430], Wilcoxon Rank Sum Test, **Figure 3D**). This pattern was still present when considering only combinations of SNVs that disrupted two separate TFBSs (0.309, [0.203-0.329]). This observation could be due to TFBS cooperativity – i.e., if one of the TFBSs is disrupted, the effect on total expression reflects reduced activity across both TFBSs. Pos-Pos pairs (-0.098, [-0.156,0.085]) and Pos-Neg pairs (-0.171, [-0.195,0.152]) behaved more additively but still demonstrated an apparent transcriptional ceiling effect (**Supplemental Figure S4D**). Combining the two most positive single variants led to a 40% increase in transcription compared to the WT sequence.

ChromBPNet models were also used to predict open chromatin activity for paired SNVs. These simulated read counts were compared to appropriate controls for each group to calculate predicted LFC that could be compared to the MPRA data. The different ChromBPNet models were concordant (mean pairwise *r =* 0.885 (95%CI = [0.874,0.896], p < 0.0001, **Supplemental Figure S4E**). As with the promoter SNVs, there was no statistical difference in median total absolute residual between ATAC-seq-trained models and DNAse-trained models (0.566 vs 0.463, p = 0.42, Wilcoxon test). The ChromBPNet LFCs for the paired SNVs were compared to the sum of the ChromBPNet LFCs for the two component SNVs. ChromBPNet predicted a greater extent of additive behavior than was experimentally determined in the MPRA (*r* = 0.95, p < 0.0001, **Figure 3E**). While ChromBPNet correctly predicted the direction of epistasis seen in the MPRA, it underestimated the magnitude of that epistasis (**Figure 3F**).

Overall, the ChromBPNet paired SNV predicted LFCs correlated with the MPRA LFCs (*r* = 0.84, p<0.0001, Pearson correlation, **Figure 3G**). However, correlations were weaker when compared for individual pair types (Pos-Pos: *r* = 0.13, p=0.08; Neg-Neg: *r* = 0.28, p<0.0001; Pos-Neg: *r* = 0.64, p<0.0001).

### Modular hybrid promoter elements increase *MYBPC3* promoter transcription

In the prior experiments, we identified combinations of SNVs capable of enhancing *MYBPC3* expression. Installation of such edits through gene editing could potentially be used to rescue *MYBPC3* haploinsufficiency as a treatment approach. To further develop this concept, we also sought to identify strategies that would allow single element insertions. TFBSs exhibit modular behavior to an extent, although sequence context and cooperativity also influence activity.^20,21^ We first determined regional effects of *MYBPC3* promoter elements that may not be obvious with the SNV saturation mutagenesis approach. We divided the *MYBPC3* promoter into eight elements based on distribution of annotated cardiac-relevant TFBSs and regions of high evolutionary conservation (referred to as M1-M8, **Figure 3H**). We then performed MPRAs using variant promoters with each possible combination of one to seven elements removed and replaced with randomized sequence (e.g., M1 removed, M2 removed, M1 and M2 removed, M4, M7, M8 removed, etc.). The MPRA also included variant promoters with each possible pair of elements swapped (**Supplemental Figure S4G**). These modifications had a range of effects on transcriptional activity in the MPRA compared to the wild-type *MYBPC3* promoter (**Supplemental Figure S6B**). Most of these modifications decreased transcription. Removal of M4 and M5 were most damaging to transcription (**Figure 3J**). These elements contain the MEF2D and GATA4 binding sites, respectively, corroborating our conclusions from the saturation mutagenesis MPRA that these regions are critical to the promoter’s function. ChromBPNet tended to underpredict the impact of these element dropouts but correctly identified M4 as a key region (**Figure 3K**).

The only variant promoter that significantly increased expression consisted of a swap between M7 and M2 (**Supplemental Figure S6A**). Since M7 contains the RXRA repressor binding site identified above, these results suggest that its transcriptional repressor function is location dependent.

With this knowledge of the general architecture of the *MYBPC3* promoter, we next tested whether insertion of hybrid promoter elements within the *MYBPC3* promoter could be capable of further increasing expression. For these experiments, we selected the *TNNT2* promoter since *TNNT2* (encoding cardiac troponin T) is a highly expressed, cardiac-specific gene, whose promoter has been previously characterized.^22^ As with the *MYBPC3* promoter, we partitioned the *TNNT2* promoter into distinct elements based on annotated TFBSs and prior studies (referred to as T1-T10, **Figure 3I**). We then tested 68 rationally designed *MYBPC3* promoter hybrid variants created by removing non-conserved regions of the *MYBPC3* promoter and inserting elements from the *TNNT2* promoter or duplicate sections of the *MYBPC3* promoter (**Supplemental Figure S6B, Supplemental Table S5-S6**). The hybrid promoters with the highest transcription levels included insertion of T4 (containing HAND1, MyoD, and YY1 TFBSs), T5 (MCAT1) or T6 MCAT2 and HSF1) either between the GATA4 and SRF/TATA sites in the *MYBPC3* promoter or within M3 of the *MYBPC3* promoter (**Figure 3L**). ChromBPNet predictions for these hybrid variants were moderately correlated with MPRA results (*r* = 0.58, p < 0.0001, **Supplemental Figure S4F, S4H**). However, this correlation was primarily driven by variants that decreased expression in the MPRA compared to WT (negative MPRA variants: *r* = 0.63, 95%CI = [0.37,0.80], p < 0.0001. positive MPRA variants: *r* = -0.18, 95%CI = [-0.55,0.26], p = 0.43. Fisher’s Z-test for difference in correlations: Z = 3.1, p = 0.0016). Nonetheless, ChromBPNet correctly predicted several of the top performing hybrid variants (**Figure 3M**).

### Rationally designed heterologous TFBS insertions increase *MYBPC3* promoter transcription

We next leveraged the MPRA approach to identify TFBS additions, combinations, and spacings that increase *MYBPC3* promoter activity. For these experiments, we used a modified version of the *MYBPC3* core promoter sequence containing the critical regulatory elements defined above to preserve regulatory context. We inserted heterologous TFBS combinations centered at the TSS -343 position, avoiding critical TFBS regions as determined above. Heterologous sequences were inserted within an otherwise transcriptionally null background sequence to parse the specific effects of TFBS grammar within the *MYBPC3* promoter context (**Figure 4A**).^23^ For these analyses, we included TFBSs of highly expressed cardiac transcription factors, as prioritized by relative expression levels from human heart RNA-seq (e.g., *MEF2D*, *SRF*, *NKX2-5*, *GATA4*, *NFIC*, *JUN*). We first tested whether single and homotypic repeats of TFBSs would increase transcription. Only a single addition of an NKX2-5 TFBS significantly increased transcription at this position (**Figure 4B**). Homotypic repeats for all of these TFBSs did not increase transcription, and, in some cases, even reduced transcriptional activity (**Figure 4B**). In contrast, multiple combinations of heterotypic TFBS insertions significantly increased transcription (**Figure 4C, Supplemental Figure S6C**). The highest level of expression was attained for an insertion of the combination of TFBSs for MEF2D, SRF, and NKX2-5 (59.8±8.9% increase compared to background sequence). Sequences and transcriptional effects for TFBS combinations are shown in **Supplemental Table S7 and S8**. ChromBPNet predictions of LFC_Background_ were generated for the synthetic CRE sequences. The ChromBPNet models predicted discrepant effects of copy number for each TF compared to the experimentally determined MPRA data, i.e., no effect for GATA4 and NKX2-5, but increased effects for MEF2D and SRF in proportion to copy number (**Figure 4D**). More synthetic CREs with heterotypic clusters were predicted to increase transcription compared to the null sequence than did so in the MPRA (**Figure 4E, Supplemental Figure S6D**). The sequence predicted to be strongest was the triple MEF2D cluster, which was observed to have an LFC_Background_ not significantly different than 0 in the MPRA. Overall, ChromBPNet predictions moderately correlated with the MPRA data (*r*=0.48, p<0.0001, **Figure 4F**). However, its predictions for the homotypic TFBS clusters were not correlated (*r*=0.09, p=0.77, **Figure 4G**). We next tested the effects of different spacings between both homotypic and heterotypic TFBS combinations (**Figure 4H-I, Supplemental Figure S7A**). Transcriptional activity was generally most increased at spacings of 10 or 50 bp. ChromBPNet predictions for these differently spaced TFBS pairs showed similar patterns and were moderately correlated with the MPRA data (*r*=0.64, p<0.0001), Figure 4J-K).

**Figure 4:**
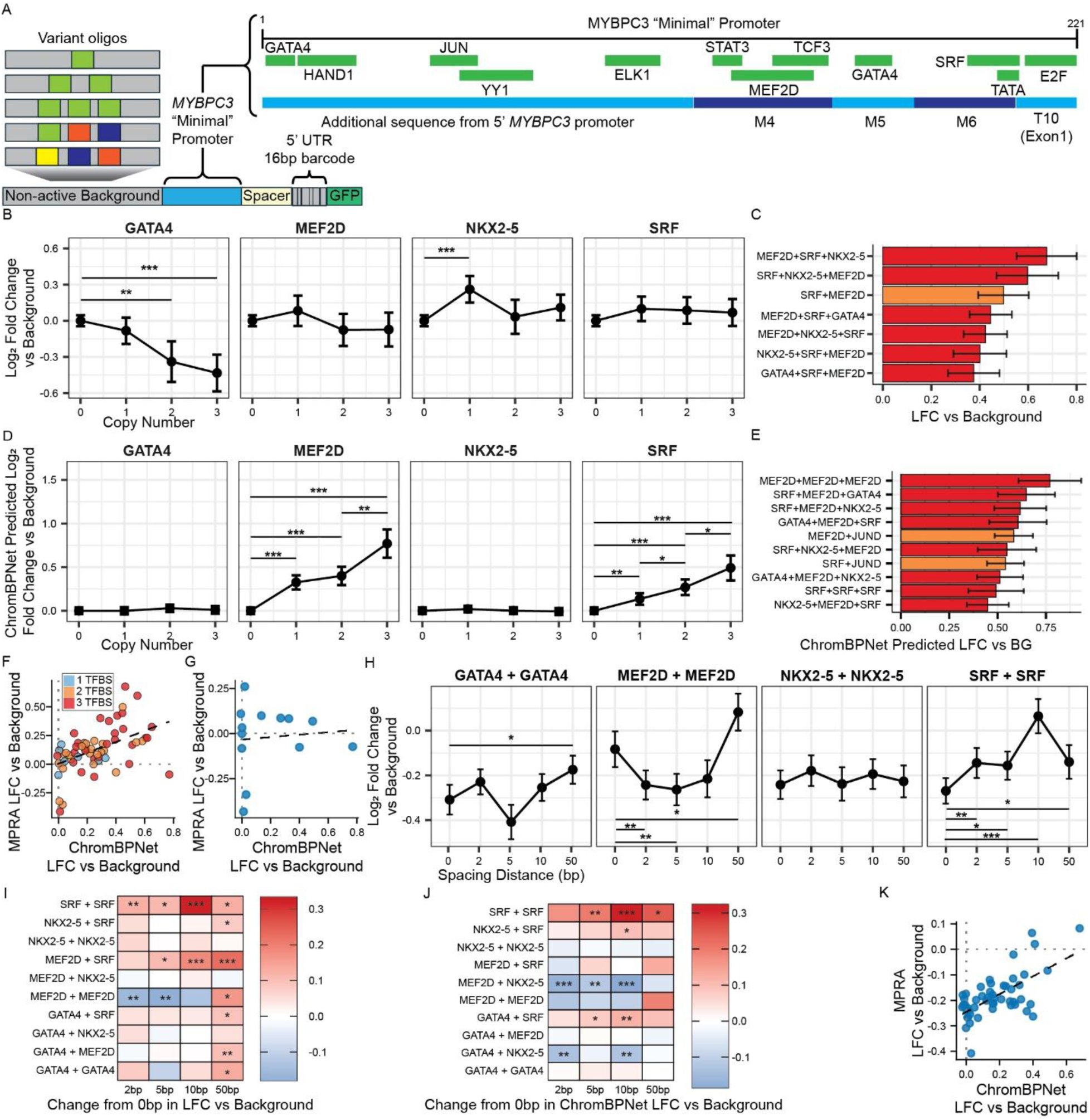
MPRA with synthetic CREs investigates regulatory grammar. **(A)** Synthetic CREs consisting of 1-3 TFBS motifs were inserted within a null background sequence upstream of a “minimal” *MYBPC3* promoter and measured using MPRA. Log_2_ fold-change compared to the null background sequence was calculated for each synthetic CRE (LFC_Background_). **(B)** Clusters of 1-3 homotypic TFBSs spaced 5bp apart were tested to determine the effect of copy number on expression for GATA4, MEF2D, NKX2-5, and SRF. **(C)** These homotypic TFBS clusters as well as multiple heterotypic TFBS combinations were tested for level of transcriptional activation in the minimal *MYBPC3* promoter context. Combinations with significant transcriptional upregulation are shown. See **Supplemental Figure S6C** for all clusters tested. **(D)** ChromBPNet models predicted the LFC_Background_ for the same clusters of 1-3 homotypic TFBSs spaced 5bp apart. **(E)** ChromBPNet models predicted the LFC_Background_ for the same homo- and heterotypic TFBS clusters. The 10 most positive are shown here. See **Supplemental Figure S6D** for all clusters tested. **(F)** Pearson correlation of MPRA data and ChromBPNet predictions for the TFBS clusters. (R=0.48, p<0.0001) **(G)** Pearson correlation of MPRA data and ChromBPNet predictions for homotypic clusters only. (R=0.09, p=0.77) **(H)** Effect of spacing between two TFBSs within the null sequence for homotypic and heterotypic pairs was assessed in the context of the minimal *MYBPC3* promoter. Spacing intervals of 0, 2, 10, and 50 bp were measured, and selected pairs shown. See **Supplemental Figure S6D** for all TFBS pairs tested. **(I)** Heatmap of spacing effects on MPRA activity for pairs of TFBSs. Color shows effect size compared to 0bp. **(J)** ChromBPNet predictions of the effects of TFBS spacing interval on LFC_Background_. **(K)** Pearson correlation of MPRA TFBS spacing data and ChromBPNet predictions for the same. (R=0.64, p<0.0001) Statistical significance: *p<0.05, **p<0.01, ***p<0.001, Wilcoxon test with Benjamini-Hochberg correction. Mean LFC_Background_ and 95%CI calculated across all replicates and barcodes for each variant for MPRA data, and calculated across the 25 models for ChromBPNet.

Altogether, these results demonstrate the capacity of rationally designed and empirically confirmed heterologous TFBS insertion to upregulate *MYBPC3* transcription. Our results also highlight how machine learning models may facilitate iterative design of synthetic CREs but perform best for smaller extents of sequence divergence.

## Discussion

Single allele loss of function variants that cause reduced protein levels (i.e., haploinsufficiency) are a common cause of disease in humans. At least 460 genes, and likely more, are associated with haploinsufficiency, accounting for a broad variety of human disorders. HCM due to *MYBPC3* loss of function variants is a prototype of human haploinsufficient disease. In this study, we have systematically interrogated the *MYBPC3* promoter to 1) create a dense map of clinically actionable, loss-of-function noncoding variant effects, 2) parse the essential TFBS grammar that regulates *MYBPC3* expression, and 3) identify promoter sequence edits that increase *MYBPC3* expression from systematic screens of thousands of variant combinations.

While gene editing approaches have begun to translate to the clinic, major hurdles limit broad utilization. Base editing to correct pathogenic variants has specifically been validated in vivo for cardiomyopathy – for example, base editing to correct an HCM-associated variant in *MYH7*.^24^ However, as is typical for most haploinsufficiency-associated genes, hundreds of unique loss of function pathogenic variants have been reported for *MYBPC3*. Creating gene editing approaches to permanently correct each of these variants would require extensive validation and safety testing to determine the optimal reagents for each individual variant. An alternative approach that still addresses the fundamental genetic etiology would be to develop strategies that restore normal levels of *MYBPC3* expression. Such strategies could include either increasing production (e.g., gene replacement therapy or upregulation of transcription) or reducing degradation (e.g., inhibiting degradation pathways). Gene replacement therapy for *MYBPC3* has shown promise in mice but will require chronic and high-level expression to be effective in humans. CRISPRa is an alternative variant-agnostic approach to upregulate gene expression and has shown potential for several haploinsufficient conditions. Notably, CRISPRa circumvents the packaging limits for current gene replacement viral vectors, and we recently demonstrated efficacy of CRISPRa as a potential rescue strategy for the large gene *DSP*. Targeting regulatory sequence elements is another promising approach with broad potential for haploinsufficient disorders. This latter approach has the potential advantage of a one-time treatment to restore expression level by up-regulation of wild-type allelic expression.

Our work here furthers this concept, utilizing MPRAs to extensively test three alternative types of sequence edits to re-calibrate *MYBPC3* expression level: 1) combinations of SNVs, 2) modular promoter element insertions, and 3) heterologous TFBS insertions. Each of these approaches was sufficient to drive meaningful increases in *MYBPC3* expression. In the case of SNV edits, iterative MPRAs allowed us to efficiently identify SNVs with significant effects, then prioritize and test combinations of these variants. For the *MYBPC3* promoter, SNV combinations demonstrated largely additive effects, enabling a relatively large dynamic range of promoter modulation. Hybrid promoter elements, systematically screened from a highly active and specific cardiac promoter (*TNNT2*), were also effective in up-regulating *MYBPC3* expression. Finally, from systematic screening of heterologous TFBS insertions, we identified combinations with substantial capacity to up-regulate *MYBPC3*. Together, these approaches form a blueprint that could be broadly implemented across other haploinsufficiency associated genes.

Our saturation mutagenesis approach for the *MYBPC3* promoter also generated a detailed map of variant effects for this noncoding region. As whole genome sequencing becomes increasingly utilized in clinical practice, this type of noncoding variant functional data will be essential for accurate interpretation of noncoding variant effects. A high degree of reproducibility across replicates and cell lines indicates that the MPRA approach can successfully map and quantify these effects.

We observed an overall fair correlation between our experimental MPRA results and ML predictions of variant effects for SNVs. Notably, correlations for SNVs affecting some TFBSs were more accurate than others, which could be influenced by local sequence context effects not fully captured in ML models. ML models also underestimated the magnitude of epistatic effect observed in pairs of SNVs. ML sequence predictions diverged markedly from experimental data for large sequence perturbations, consistent with a prior study that reported inaccurate ML predictions for sequences that highly diverge from naturally occurring sequences.^25^ For example, ML models predicted completely additive behavior for homotypic repeat TFBS insertions, but MPRAs revealed a clear plateau effect. In contrast, specific heterotypic combinations were able to robustly increase transcription. These comparisons highlight the relative strengths of ML models for SNV predictions but limitations in predicting effects of large sequence perturbations.

In conclusion, we have mapped the *MYBPC3* promoter with base pair precision, revealing the TFBS grammar essential for driving expression of this gene. Further, we have systematically screened thousands of promoter sequence edits to identify variant combinations that increase transcription. These approaches may be broadly useful to develop variant-agnostic treatments for diseases caused by haploinsufficiency.

## Methods

### Culture and differentiation of induced pluripotent stem cell-derived cardiomyocytes (iPSC-CMs)

Control iPSCs (WTC, Coriell) and a previously established healthy female line^26,27^ were cultured in mTeSR-plus (Stem Cell Technologies). Cardiomyocyte differentiation via Wnt modulation and subsequent purification in glucose-deprived, CDM3-lactate media was performed as previously described.^26,28,29^ Purified hPSC-CMs were replated as monolayers, cultured in CDM3-lactate media for an additional 4 days followed by oxidative phosphorylation media for an additional 4 days, to promote maturation and halt cell replication, as described.

### Lentivirus-based massively parallel reporter assays

Lentiviral MPRA libraries were constructed based on prior protocols with modifications to facilitate promoter mapping.^14^ We first modified the pLS-SceI plasmid backbone (Addgene #137725) to remove native BbsI restriction sites (T4200A and A5285T) through digestion with BbsI and replacement with a corrected gene fragment using Gibson assembly. We then digested this backbone with SbfI and AgeI to clone in a modular CRE-reporter sequence containing separate Golden Gate insertion sites for CRE (flanking BsmBI sites) and barcode (flanking BbsI sites) insertions (synthesized from Twist Biosciences with Gibson assembly adaptors on each end). We used 3 versions of the CRE-reporter sequence for different purposes: 1) 5’ constant half of the *MYBPC3* promoter with BsmBI insertion site placed to map the 3’ half, 2) 3’ constant half of the promoter with Bsm’BI sites placed to map the 5’ half, and 3) shortened 221bp version of the *MYPBC3* promoter to test upstream CRE insertion (See **Supplemental Table S9** for sequences). An upstream *ACTC1* enhancer was also included in the CRE-reporter to increase overall transcription. A subset of MPRAs were also performed without the *ACTC1* enhancer.

We designed 300 bp oligo pools comprised of 250 bp promoter or CRE variant sequences with 25 bp flanking adaptors; sequences were then synthesized (Twist Biosciences). See **Supplemental Table S10** for oligo sequences. Adaptor sequences were used as PCR primers to amplify each oligo pool from the synthesized fragments (**Supplemental Table S11**). Each amplified pool was then cloned into respective BsmBI cloning sites in the CRE-reporter modules using Gibson assembly (NEBuilder, New England Biolabs). Resultant plasmid pools were electroporated into 10-beta electrocompetent *E. coli* (New England Biolabs) and grown in overnight 50 ml liquid cultures. Serial dilutions of transformed bacteria were concurrently used to estimate total colony forming units (CFUs) – at least 10 CFUs per oligo variant were required to ensure library diversity. Next, this process was repeated to insert a 16bp randomized barcode fragment (Integrated DNA Technologies) in place of the BbsI dropout site to create the final barcoded CRE variant plasmid pool, requiring at least 115 CFUs per oligo variant for an estimated coverage of 95% edits with at least 100X coverage. CRE-reporter sequences were amplified from the plasmid pool for high-throughput long-read sequencing (University of Michigan Advanced Genomics Core) to associate barcodes with promoter/CRE variants. A mean coverage of at least 25X unique barcodes per variant in the library was required to advance to lentiviral production. Lentivirus was prepared from this plasmid pool by the University of Michigan Vector Core. Titers were determined by GFP expression with fluorescence microscopy.

Each oligo library was transduced in at least 3 separate rounds of transductions. Transductions were performed in iPSC-CMs differentiated from each of the different iPSC lines above. Each round of transduction was performed in >1,000-fold iPSC-CMs per variant in the oligo library to ensure both coverage of all variants and many replicate measurements per variant. To avoid iPSC-CM differentiation batch effects, iPSC-CMs were combined from multiple cryopreserved differentiations simultaneously for each transduction round. iPSCs-CMs were recovered from thawing and cultured for five days prior to transductions. After three additional days, DNA and RNA were extracted simultaneously (Allprep DNA/RNA Extraction Kit, Qiagen). RNA was reverse transcribed (Omniscript RT Kit, Qiagen). Libraries were prepared by PCR amplification for Illumina sequencing (primers in **Supplemental Table S11**). Both DNA and cDNA were then Illumina sequenced to cover the barcode region (University of Michigan Advanced Genomics Core).

### Barcode count analysis

Long-read sequencing data for associating barcodes with variants were processed using samtools v1.17, cutadapt v4.4, and a python script adapted from Cooper et. al., 2022 to generate a variant-to-barcode dictionary. Illumina sequencing data used to count barcodes for experimental replicates were processed using cutadapt v4.4 and a python script adapted from Cooper et. al., 2022^30^. Barcode counts for both RNA and DNA were then analyzed in R v4.4.1^31^. Only barcodes with at least 10 DNA and at least 10 RNA counts were included. Barcodes within each replicate were also removed if their RNA count /DNA count ratios were outside Q1-Q3 ± 1.5*IQR for a particular variant. Additionally, to average out potential confounding of expression measurements due to specific barcode sequences or lentiviral insertion sites, only variants associated with at least 10 unique barcodes and represented in at least two replicates were included. We performed several downstream analyses with sets of oligos that were tested in the same experiment. After quality filtering, raw counts were aggregated over all barcodes for a given variant and used as input for downstream testing. mpralm^32^ (version 1.26.2) was used to normalize aggregated counts and fit linear models to estimate the log_2_(fold-change) in activity between allele pairs (test variants and their paired control). To rank allele pairs by significance, empirical Bayes moderated t-statistics were computed relative to a fold-change threshold of 0. For analysis of paired SNVs and TFBS insertions, which did not fit model assumptions of mpralm, expression effects were calculated as mean log_2_ fold-change compared to an appropriate control across all replicates on a barcode-by-barcode basis. The means for each variant were then compared using Wilcoxon rank sum test with Benjamini-Hochberg correction.

### TFBS motif analysis

Position weight matrices (PWM) were generated from MPRA data for 31bp windows centered on each SNV. These were matched to canonical motifs from JASPAR 2020^17^ using motifmatchr and TFBSTools^33^ in R. For each SNV, a delta score was calculated as the difference in PWM log-likelihood scores between the alternate and reference alleles. Negative delta scores indicate motif disruption and positive delta scores indicate strengthening or creation of a TFBS motif.

To generate empirical logo-style TFBS motif plots, base importance was calculated using MPRA data. For each position and variant, impact score was calculated as LFC_Variant_ / LFC_KO_ where LFC_KO_ is a position-specific knockout baseline estimated from mean rolling triplet inversion disruption of that site or, for variants from -428 to -219, estimated from a global mean triplet effect ratio applied to the mean SNV effect in that TFBS. Tolerance values were normalized to probabilities p_b_ over bases, and Shannon information content was calculated as IC = max[0,2+∑_b_p_b_log_2_(p_b_)].^34,35^ The displayed letter height for each base was p_b_ × IC, yielding a constraint-weighted logo comparable in scale to canonical bits. These MPRA-derived logos were then compared to canonical motifs using Pearson correlation.

### ChromBPNet Analysis

*In silico* mutagenesis (ISM) analyses were performed for the MPRA variants using 25 ChromBPNet models trained on ATAC-seq and DNAse-seq data from 25 left heart ventricle tissue samples from the ENCODE portal^12^. For each variant, a 2,114 bp window centered on the region of interest was extracted from the reference genome and passed to each model. For wild-type, null control, and each variant sequence, the model predicted ATAC-seq read count, averaged across five cross-validation folds. These predicted read counts were used to calculate log_2_ fold-change compared to appropriate controls.

### AlphaGenome Analysis

ISM analyses were also performed for promoter SNVs using AlphaGenome using an automated Python (v3.11) workflow (https://github.com/Kaploc23/AlphaGenome). The same promoter region of *MYBPC3* was assessed with a surrounding genomic sequence context of 150 kb. Mutagenesis at each position was evaluated against CAGE, ATAC-seq, DNase-seq, and ChIP-seq tracks. Requests were conditioned on ontology-based filtering for left ventricle.

### Paired SNV Analysis

To evaluate the additive or epistatic interactions of paired promoter SNVs, we employed a weighted least squares linear regression using the base lm function in R. We fit a model to replicate-level MPRA data for each pair using a statistical weight for each replicate based on the variance and number of unique barcodes represented by that replicate (specifically, the inverse variance of its log_2_ mean). The model was fit to LFC_Pair_ ∼ LFC_SNV1_ * LFC_SNV2_, with the coefficient of the interaction term representing the quantitative synergy score. A synergy score of 0 represents perfect additive effect. The associated p-values were corrected using the Benjamini-Hochberg correction with an FDR of 0.05.

## Disclosures

ASH has consulted for Tenaya Therapeutics, Lexeo Therapeutics, Preload Therapeutics, Cytokinetics, and Alexion Pharmaceuticals. JSW has received research support from Bristol Myers Squibb, has acted as a paid advisor to Health Lumen, Tenaya Therapeutics, Solid Biosciences, Hopkins Van Mil on behalf of Genomics England, and HEOR Ltd on behalf of Rocket Pharmaceuticals, and is a founder with equity in Saturnus Bio.

## Acknowledgments

Elements of Figure 1A were produced in Biorender. We thank Stanford University and the Stanford Research Computing Center for providing computational resources and support as part of the Sherlock High-Performance Computer Cluster.

## Sources of Funding

Research reported in this publication was supported by the National Institutes of Health under Award Numbers R01-HL171074 (ASH), T32 HL007853 (EMS), K12 HD028820 (JKM), K08HL179262 (JKM), R35GM153286 (JK), 25PRE1372132 (AR), T32GM156550 (AR), T32GM145470 (AR), R01HL171609 (JME), R01HL184893 (JME), Medical Research Council (UK), British Heart Foundation [RE/24/130023], the NIHR Imperial Biomedical Research Centre, Sir Jules Thorn Charitable Trust [21JTA], and the Wellcome Trust [226083/Z/22/Z]. JSW is supported by CureHeart, the British Heart Foundation’s Big Beat Challenge award [BBC/F/21/220106]. JME acknowledges support from the Applebaum Foundation and the BASE Research Initiative at Lucile Packard Children’s Hospital. MUS was supported by the NSF Graduate Research Fellowship (DGE-1656518) and a graduate fellowship award from Knight-Hennessy Scholars at Stanford University. Content is solely the responsibility of the authors and does not necessarily represent the official views of the funders.

For the purpose of open access, the authors have applied a Creative Commons Attribution (CC BY) license to any Author Accepted Manuscript version arising.

## Supplemental Figures

**Figure S1:**
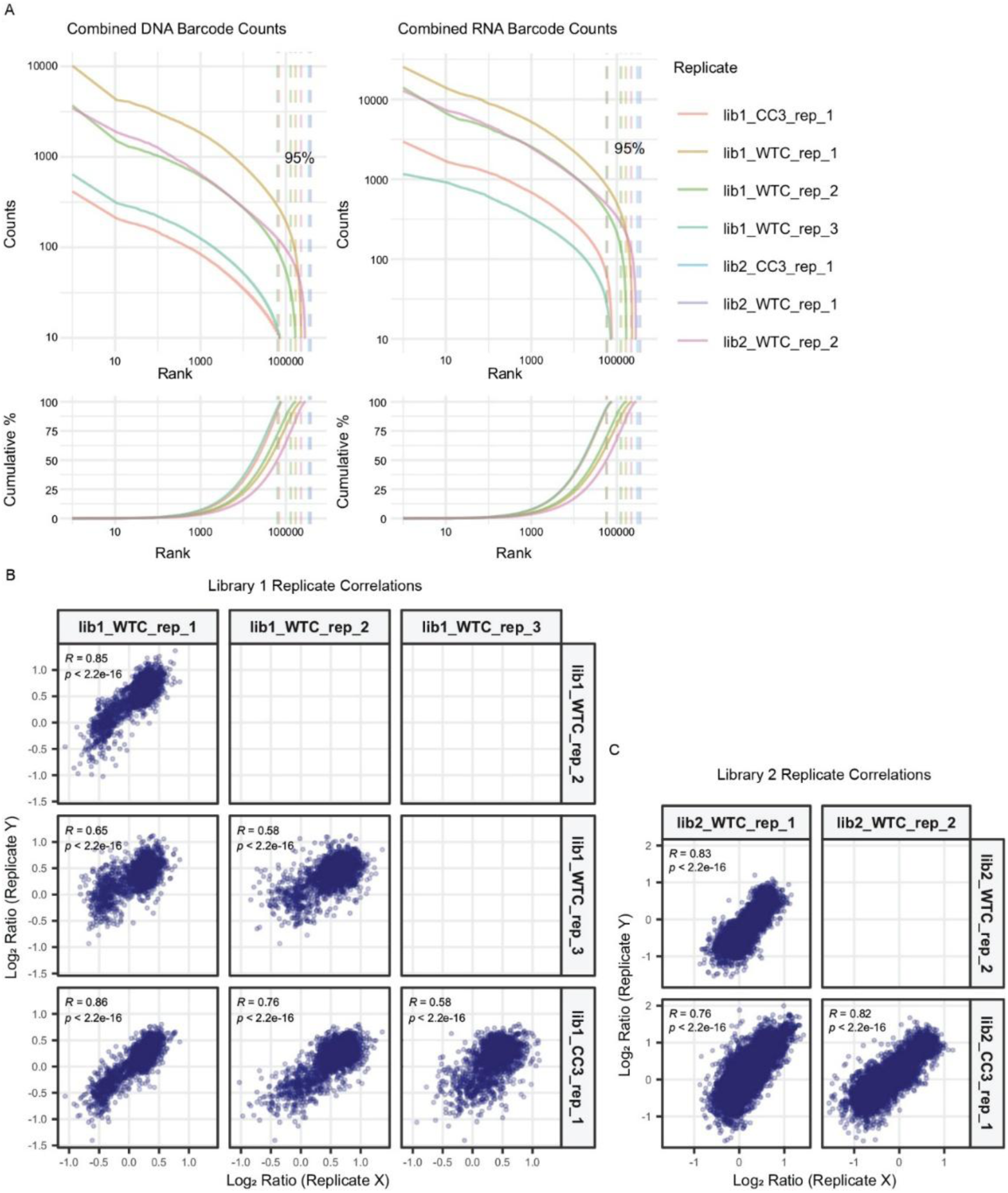
Waterfall plots and replicate correlations for MPRA. The MPRA library 1 consisted of SNVs for *MYBPC3* promoter positions 211-456, as well as rolling triplet variants and *TNNT2*-*MYBPC3* hybrid promoters. Library 2 consisted of SNVs for *MYBPC3* promoter positions 1-210, combinations of SNVs in positions 211-456, as well as *MYBPC3* hybrid promoters and synthetic enhancers. **(A)** 95% of barcodes had high DNA and RNA counts. Barcodes with fewer than 10 counts in a given replicate were excluded from analysis. Library 1 **(B)** and library 2 **(C)** Pearson correlations demonstrate high concordance of variant mean DNA/RNA ratio data across biological replicates.

**Figure S2:**
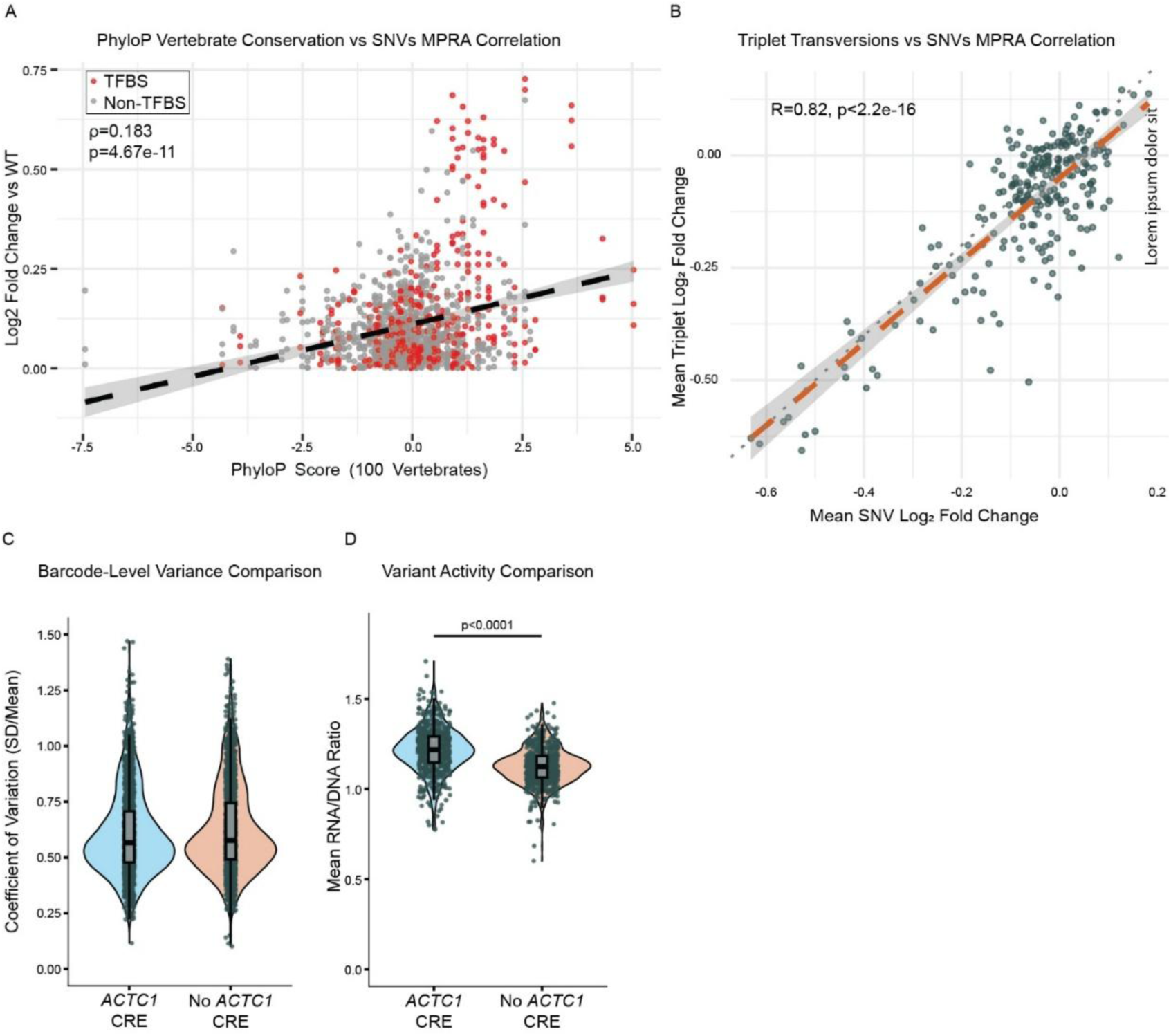
Analysis of alternative approaches to MPRA library construction. **(A)** Vertebrate conservation is only very weakly correlated with SNV MPRA effect size (Spearman correlation). **(B)** Rolling triplet transversions represent an alternative to saturation mutagenesis for TFBS mapping through MPRA, trading mapping resolution for the ability to cover the same genomic territory with decreased library complexity, thus requiring fewer cells for each experimental replicate. Impact on activity for each position was highly correlated between the triple variants and SNVs. (Pearson correlation) **(C-D)** A portion of the library (SNVs for positions -428 to -219) was designed with paired duplicates lacking the upstream ACTC1 enhancer to test if that enhancer improved signal:noise in the MPRA. While versions with the enhancer increased mean barcode activity (mean RNA/DNA ratio = 1.22 with enhancer vs 1.13 without, p<0.0001 Wilcoxon test), it did not improve the coefficient of variation in a biologically meaningful way (CV=0.577 with enhancer vs 0.566 without, p<0.01 Wilcoxon test).

**Figure S3:**
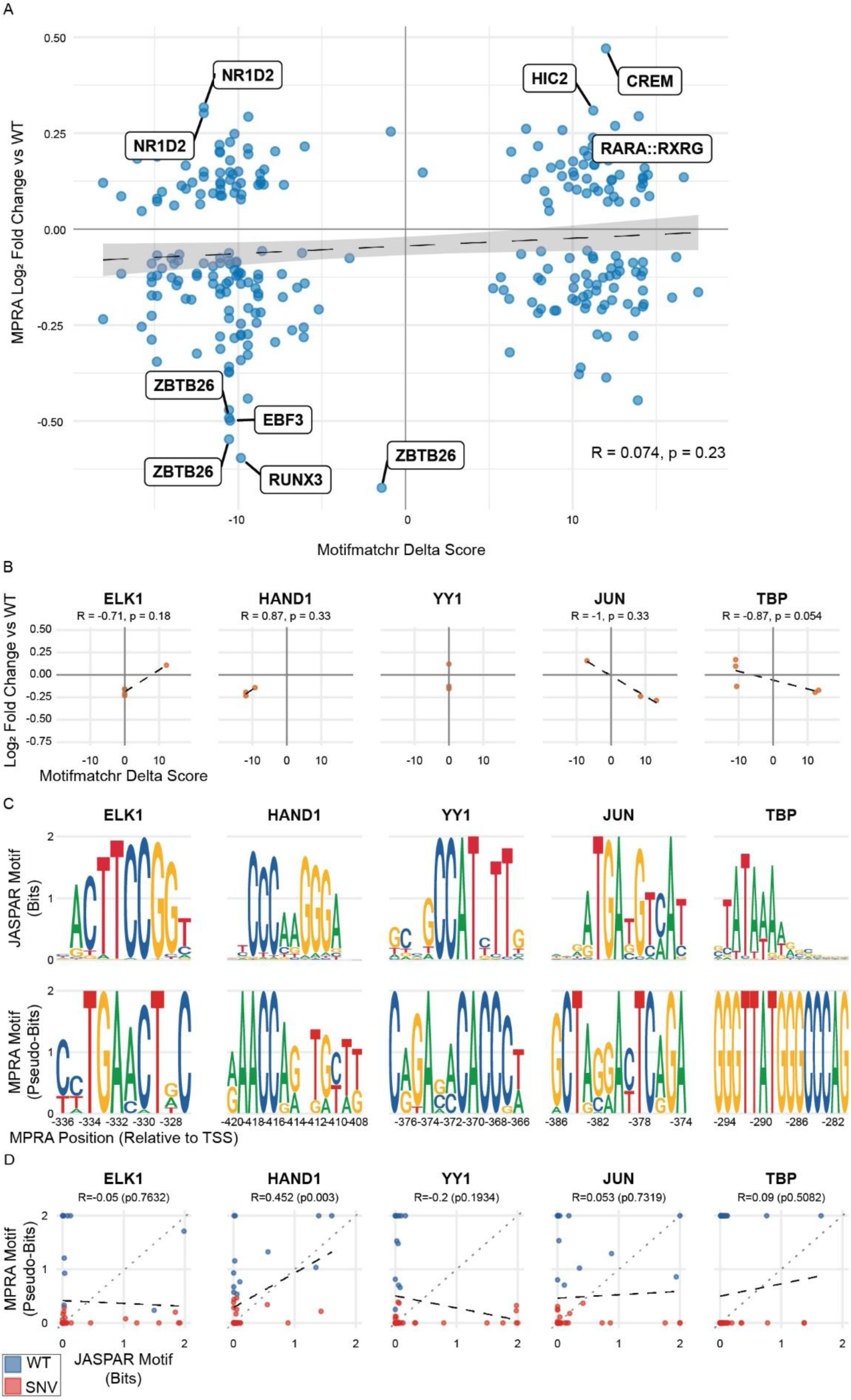
Motifmatchr analysis of non-annotated SNVs and additional annotated TFBSs. **(A)** Predicted SNV impact on TFBS binding affinity (Delta Score) was calculated using motifmatchr^16^ (see Methods) and compared to impact on transcription in the MPRA for SNVs outside of TFBS annotations (Pearson correlation). **(B)** Correlation of predicted SNV impact on binding efficiency compared to impact on MPRA transcription (Pearson correlation). Additional annotated TFBSs not shown in Figure 2C. Positive correlation indicates that TF is acting as an activator while negative correlation suggests it is acting as a repressor. **(C)** Logo-style motifs for TFBSs were generated empirically from MPRA SNV data by calculating a contribution score for each base at each position (see Methods) and are shown beneath their canonical motifs from JASPAR^17^. Additional TFBSs not shown in Figure 2D. **(D)** The contribution score for each base calculated in generating the logo-style motifs was compared to the contribution scores from the canonical JASPAR motif (Pearson correlation). Additional TFBSs not shown in Figure 2E.

**Supplemental Figure S4:**
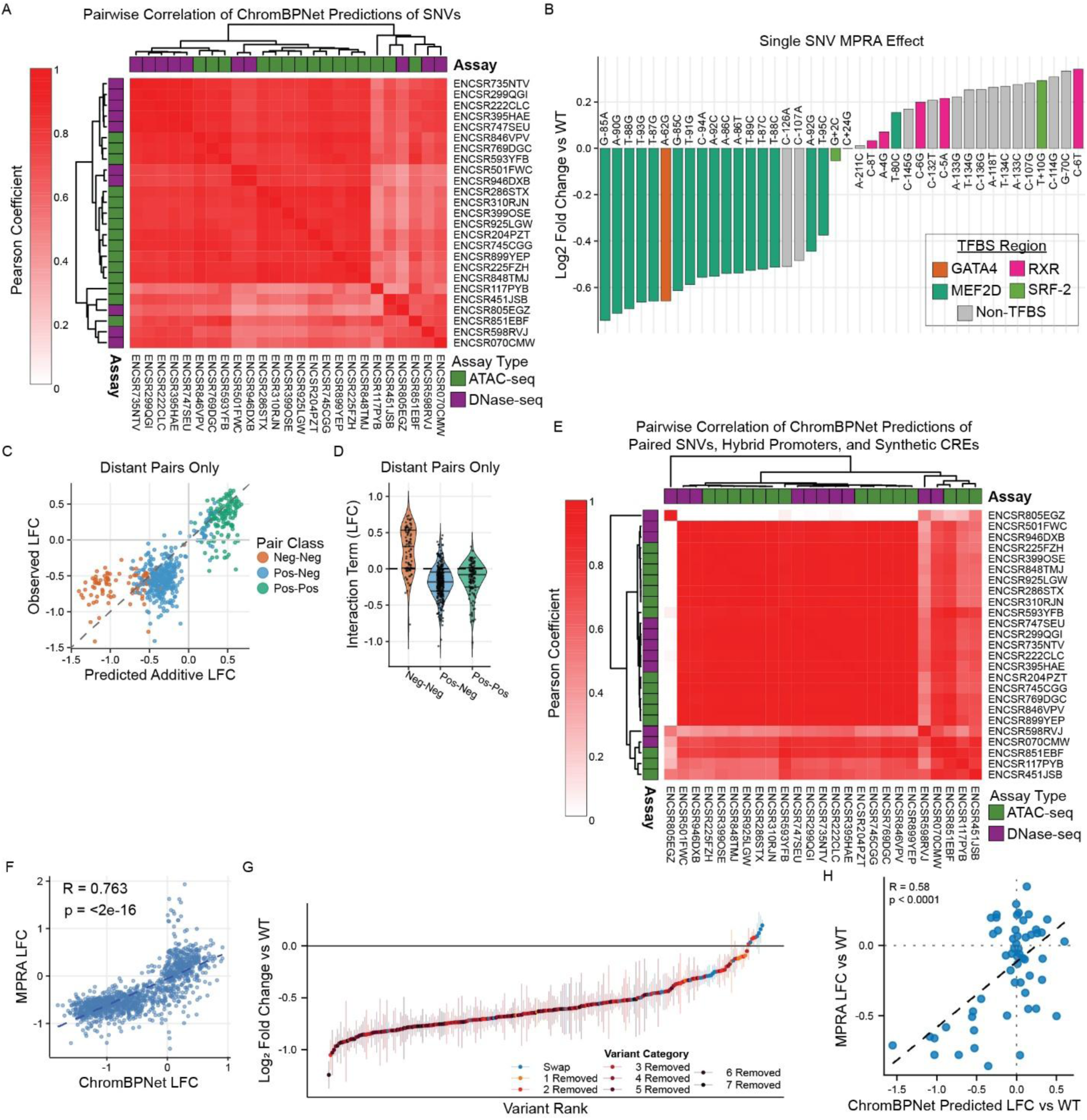
ChromBPNet model pairwise correlation, paired SNVs, and additional tested hybrid promoters. **(A)** 25 ChromBPNet models were trained on different DNase and ATAC-seq datasets from human left ventricle samples. Pairwise analysis of these models shows high correlation on the simulated ATAC-seq read counts predicted for each *MYBPC3* promoter SNV variant sequence in the MPRA libraries (mean Fisher z-transformed *r* = 0.76, 95%CI = [0.76-0.77], mean p value < 0.00001). **(B)** LFC_WT_ of the SNVs with the 20 largest positive and 20 largest negative MPRA effects that were selected for the paired SNV analysis. **(C)** A weighted linear model was created for each possible pair among these 40 variants using the single SNV LFC_WT_ to predict the paired LFC_WT_ assuming additive effects (Figure 3C). In an alternate model, excluding pairs of SNVs that are within 10bp of each other and presumably affect the same TFBS did not meaningfully change the correlation of LFC_WT_ predicted by the weighted linear model compared to observed MPRA LFC_WT_ (R = 0.80, p < 0.0001, Pearson correlation). **(D)** This alternate model has the same pattern of sub-additivity in each category seen in the fully inclusive model. **(E)** Pairwise analysis of the ChromBPNet predictions for paired SNVs (Figure 3C), hybrid *MYBPC3* promoter elements (Figure 3H), and synthetic CREs (Figure 4E). The 25 models trained were highly concordant (mean pairwise Fisher z-transformed *r* = 0.88, 95%CI = [0.87,0.89], mean p-value = 0.0098). **(F)** The ChromBPNet models were highly correlated with the MPRA data. (*r* = 0.763, p < 0.0001) **(G)** The *MYBPC3* promoter was divided into 8 elements (Figure 3H). Each combination of 1, 2, 3, 4, 5, 6, and 7 elements were removed and replaced with randomized sequence. Each element was also swapped for each other element. These variants were compared in an MPRA to the WT promoter. **(H)** ChromBPNet was moderately correlated with MPRA data for the hybrid promoters but tended to underpredict promoters that increased expression in the MPRA (*r* = 0.58, p<0.0001). Significance: *FDR<0.05, **FDR<0.01, ***FDR<0.001, Wilcoxon rank sum test with Benjamini-Hochberg correction and 95%CI unless otherwise noted.

**Supplemental Figure S5:**
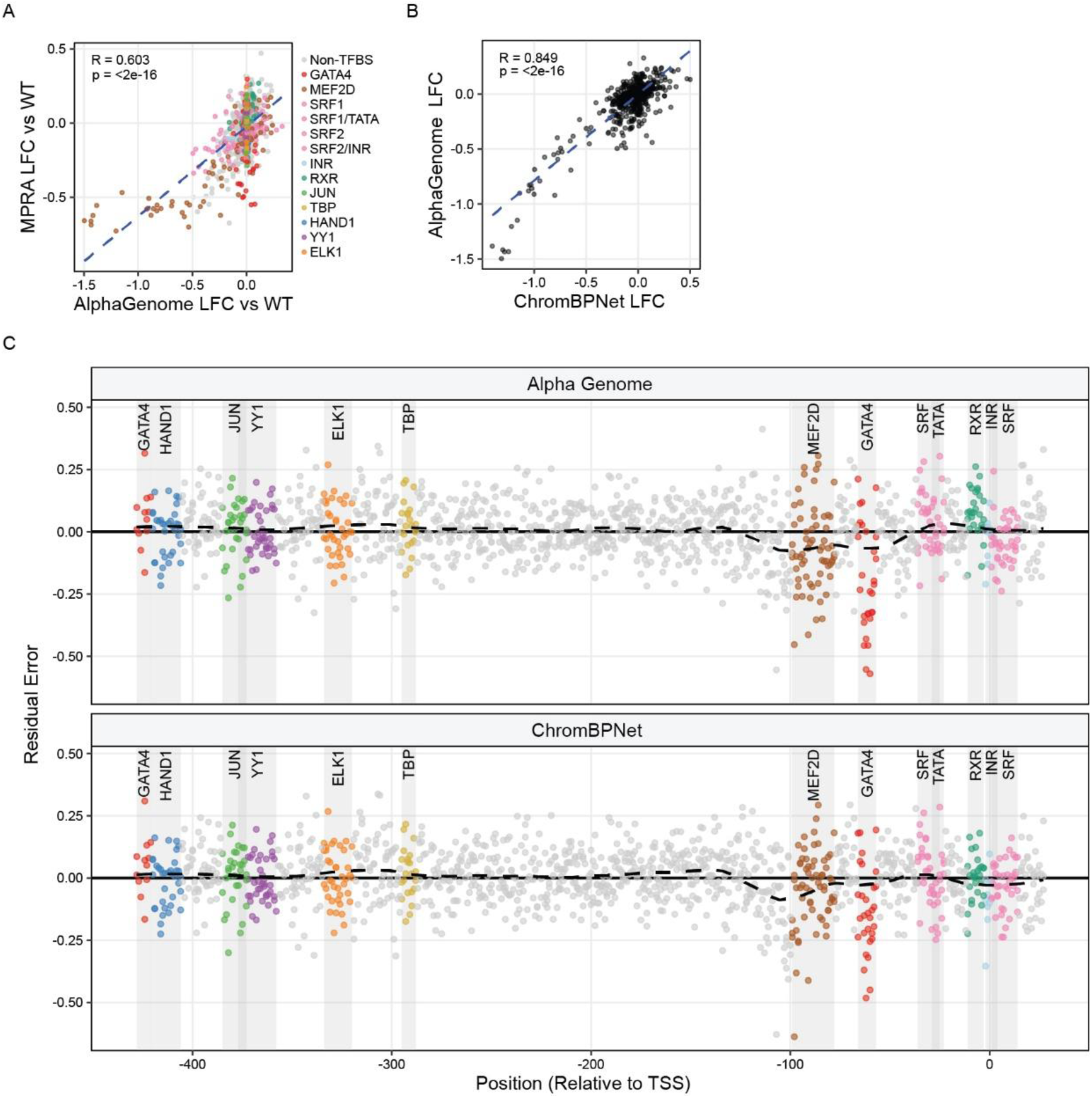
AlphaGenome predicted SNV effect on expression similarly to ChromBPNet. **(A)** AlphaGenome was used to perform in silico saturation mutagenesis of the *MYBPC3* promoter with 150kb of context. The predicted effects of SNVs on expression were moderately but significantly correlated with the MPRA experiment data (*r* = 0.60, p<0.0001). **(B)** AlphaGenome predictions were highly correlated with the ChromBPNet predictions for corresponding variants (*r* = 0.85, p<0.0001). **(C)** Analysis of the scale-adjusted residual error from the correlation of the two models with the MPRA data shows similar patterns of over and under predictions throughout the promoter sequence.

**Supplemental Figure S6:**
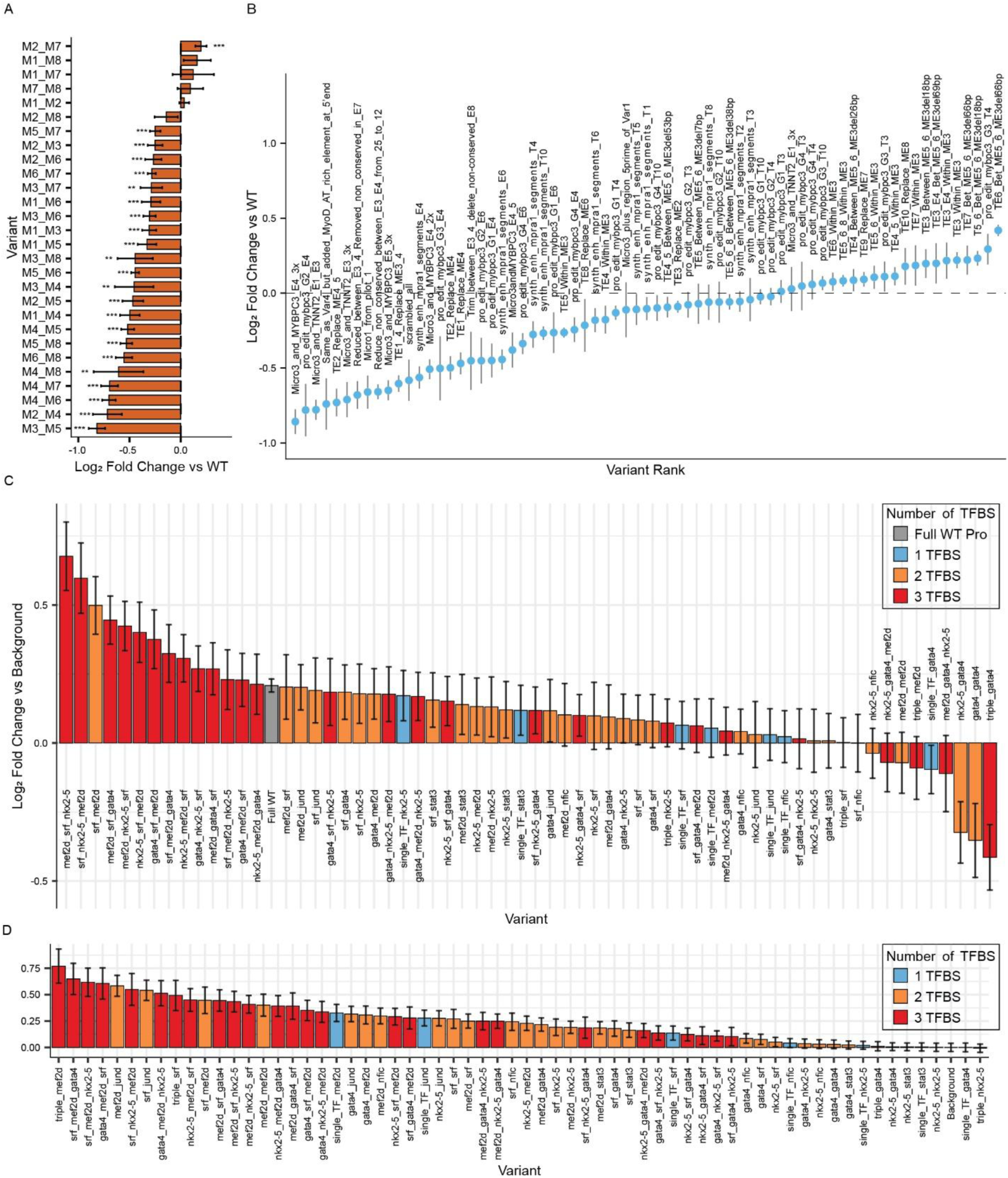
Hybrid promoters and synthetic CREs can drive transcription. **(A)** Effects of each element swap variant on transcription. **(B)** 68 *TNNT2* and *MYBPC3* hybrid promoters were tested in an MPRA and compared to the full WT *MYBPC3* promoter. **(C)** All synthetic CREs tested in the MPRA compared to the null background sequence. **(D)** ChromBPNet models predicted the LFC_Background_ for the same homo- and heterotypic TFBS clusters. Significance: *FDR<0.05, **FDR<0.01, ***FDR<0.001, Wilcoxon rank sum test with Benjamini-Hochberg correction and 95%CI unless otherwise noted. Mean LFC_Background_ and 95%CI calculated across all replicates and barcodes for each variant for MPRA data, and calculated across the 25 models for ChromBPNet.

**Supplemental Figure S7:**
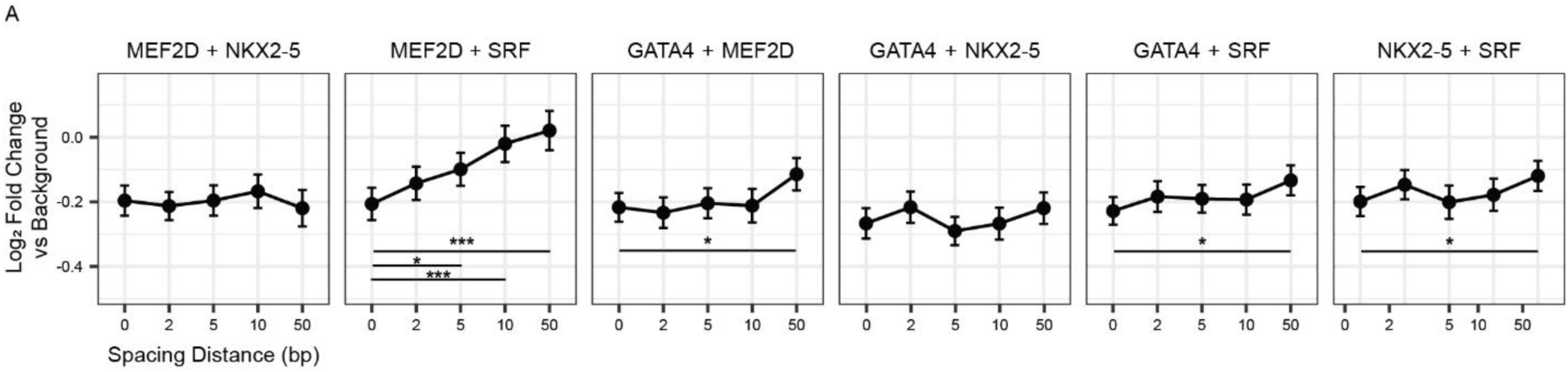
Impact of TFBS spacing on transcriptional strength. **(B)** Effect of spacing between two TFBSs within the null sequence for homotypic and heterotypic pairs was assessed in the context of the minimal *MYBPC3* promoter. Spacing intervals of 0, 2, 10, and 50 bp were measured, heterotypic pairs shown here (homotypic pairs are shown in Figure 4H).

