## Supplemental Methods for "Mapping and rewiring the *MYBPC3* promoter for rescue of haploinsufficiency driven hypertrophic cardiomyopathy"

Sequences for the MPRA plasmid backbone, custom gene fragments, and PCR primers can be found in Supplemental Tables S8 and S10. Full MPRA library oligo sequences can be found in Supplemental Table S9.

**MPRA Part 1: MPRA Plasmid Backbone Assembly**

Starting with modified pLS-SceI plasmid “modified pLS-SceI lentiMPRA plasmid backbone” which has T4200A and A5285T mutations to eliminate BbsI cut sites.

1) Linearize plasmid:

Plasmid	3ug

SbfI-HF	3ul

AgeI-HF	3ul

10X NEB cutsmart buffer	5ul

ddH2O	to 50ul total

Incubate at 37C for 1 hour.

2) Perform gel electrophoresis on 1% agarose gel at 120V for 45mins. Cut out linearized fragment.
3) Extract linearized DNA from the gel fragment using Qiaquick Gel Extraction kit (Qiagen) according to kit protocol.
4) Set up Gibson assembly reaction with linearized plasmid and backbone gene fragment using NEBuilder HiFi DNA assembly master mix (New England Biolabs). Use online NEBuilder calculator (https://nebuildercalculator.neb.com/) to determine reaction amounts for correct molar ratio. Incubate 1 hour at 50C.
5) Transform Gibson assembly product into DH5-alpha competent *E. coli* (New England Biolabs) following manufacturer protocol. Plate on ampicillin plates and incubate overnight.
6) Pick isolated colonies from the plates and grow overnight in 2ml ampicillin cultures.
7) Extract DNA from 1ml of each culture using QIAprep Spin Miniprep Kit (Qiagen) per kit protocol.
8) PCR screen colonies for insertion using NEBNext High-Fidelity 2X PCR Master Mix (New England Biolabs):

Primer Backbone_Amp_F (10uM)	0.5ul

Primer Backbone_Amp_R (10uM)	0.5ul

Plasmid DNA	1ul

ddH2O	18ul

2X NEBNext HiFi PCR Master Mix	20ul

Run: 94C 1min -> [94C 10s -> 67C 30s -> 72C 45s]x35 cycles -> 72C 2mins

9) Perform gel electrophoresis on 2% agarose gel at 100V for 60mins. Identify colonies with an amplified band.
10) Sequence plasmids with positive PCR screen to confirm insertion. If no positive screens, pick more colonies and repeat.
11) Once sequence is confirmed for a colony, use remaining 1ml of bacterial culture to inoculate a 100ml culture and incubate overnight.
12) Extract DNA from 100ml culture using Purelink HighPrep Plasmid Midiprep kit (Invitrogen) per kit protocol.

Plasmid backbone is now ready for variant oligo insertion.

**MPRA Part 2: Library Pool Amplification**

Synthesized variant oligo pool (Twist Biosciences) will come with a limited amount of DNA. This amplifies the pool so there’s a larger supply.

1) PCR 1

Snythesized Oligo pool	1ul

Pool specific F primer (100uM)	1ul

Pool specific R primer (100uM)	1ul

ddH2O	97ul

NEBNext 2x high fidelity PCR master mix	100ul

Split reaction (200ul total) into 5 PCR tubes.

98C 2mins -> [98C 15sec, Pool specific primer Tm 20s, 72C, 30s]x5 cycles -> 72C 5mins

2) Pool PCR 1 product, purify using Highprep PCR cleanup beads (MagBio) using manufacturer protocol. Elute into 50ul Buffer EB.

3) PCR 2

PCR 1 product	~100ng

Pool specific F primer (100uM)	2ul

Pool specific R primer (100uM)	2ul

ddH2O	variable to 200ul total

NEB 2x high fidelity PCR master mix	200ul

Split reaction (400ul total) into 10 PCR tubes.

98C 2mins -> [98C 15sec, Pool specific primer Tm 20s, 72C, 30s]x12 cycles -> 72C 5mins

4) Pool PCR 2 product. Add 200ul 6x loading dye. Load all of product on 1.5% agarose gel and run at 100V for an hour.

5) Cut bands of the expected size out of gel.

6) Extract DNA from the gel fragments using Qiaquick Gel Extraction kit (Qiagen) following kit protocol with these modifications: After melting the gel fragments, before adding isopropanol, combine all buffer QC/melted gel into a single 15ml tube. Vortex, add 1 volume of isopropanol, and vortex again. Put 2 columns on a vacuum manifold. With the manifold on, alternate loading 800ul of the mixture from the 15ml tube onto each of the columns. When the column drain, add another 800ul, until all of the mixture has been put through the columns. Then continue the kit as instructed, combining the eluted DNA at the end.

7) Purify the DNA extracted from the gel using Highprep PCR cleanup beads (MagBio) using manufacturer protocol. Elute into 50ul Buffer EB.

8) Run 20-30ng of the purified DNA on a 1.5% agarose gel to check quality.

9) Sequence some of the pool using amplicon sequencing (we use Plasmidsaurs) to check pool quality and diversity. If it is high quality, the pool is ready for library assembly. If assembling oligos from multiple pools into the same backbone, mix molar ratios proportional to number of oligos in each pool.

**MPRA Part 3: Plasmid pool assembly**

Starting with backbone plasmid and amplified oligo pool.

1) Digest 5ug of backbone plasmid with Esp3I (Thermo Scientific)
   a. Mix 5ug backbone, 5ul 10x Fast Digest buffer, 2.5ul Esp3I, ddH2O to 50ul total
   b. Incubate at 37C for 1 hour
2) Run on 1% agarose gel at 100V for an hour. Cut out linearized backbone.
3) Extract DNA from the gel fragments using Qiaquick Gel Extraction kit (Qiagen) following kit protocol.
4) Set up Gibson assembly reaction using NEBuilder HiFi DNA assembly master mix (New England Biolabs) with amplified pool using 0.0125 pmol linearized backbone and 0.025pmol amplified pool oligos. Also set up negative control reactions for each backbone-pool combination using ddH2O in place of oligos.
   a. Mix 0.0125pmol linearized backbone, 0.025pmol amplified pool, 10ul NEBuilder master mix, ddH2O to 20ul
   b. Mix 0.0125pmol linearized backbone, 10ul NEBuilder master mix, ddH2O to 20ul (control reaction)
   c. Incubate at 50C for 1 hour
5) Purify Gibson assembly reaction products using 0.65x volume Highprep PCR cleanup beads (MagBio) using manufacturer protocol. Elute into 44ul buffer EB.
6) Digest again with Esp3I (to digest any remaining uncut backbone).
   a. Mix 44ul Gibson assembly product, 5ul Fast Digest buffer, 1ul Esp3I
   b. Incubate at 37C for 1 hour
7) Purify digest using 1.8x volume Highprep PCR cleanup beads (MagBio) using manufacturer protocol. Elute into 20ul buffer EB.
8) Electroporate purified Gibson assembly products and controls into 10-beta electrocompetent *E. coli* (New England Biolabs) using Gene Pulser Xcell electroporation device (Biorad) with 0.1mm electroporation cuvettes (Biorad). Electroporation settings: 2000V, 200 Ohms, 25uF
   a. Prep:
      i. Prewarm stable outgrowth media to 37C
      ii. Prewarm agar plates (# electroporations x3 +1)
      iii. Pre-chill cuvettes on ice (# electroporations + 1 for positive control)
   b. Thaw 25ul electrocompetent cells on ice for each electroporation + 1 for pUC19 positive control
   c. For each Gibson assembly reaction (including negative control), and for pUC, put 4ul DNA into a microcentrifuge tube, label, and put on ice.
   d. Add 25ul cells to each tube, flick gently, put back on ice for at least 5 mins.
   e. Set up for electroporation
      i. Set one p200 pipette to 38ul
      ii. Set one p200 pipette to 150ul
      iii. Set a p1000 pipette to 975ul
      iv. Get outgrowth media from warmer
      v. Set up electroporator settings -> 2000V, 200 Ohms, 25uF
   f. Electroporate each Gibson assembly reaction and controls
      i. Use p200 and skinny tip to transfer 28ul cells/dna mix to bottom of cuvette.
      ii. Tap cuvette until all bubbles are gone.
      iii. Dry sides of the cuvette
      iv. Pulse with electroporator
      v. Immediately use p1000 to add 975ul outgrowth media and pipette to mix.
      vi. Use p1000 to transfer as much as you can to a new microcentrifuge tube.
      vii. Use the other p200 and skinny tip to transfer the remainder to the tube.
   g. Once all electroporations are done, move tubes to 37C shaking incubator for 1 hour.
   h. Plate using serial dilution. Positive control can be diluted 1:10 and plated on a single plate. For everything else:
      i. Put 50ml LB in a flask with appropriate antibiotic
      ii. Put 90ul LB on three plates labeled with plasmid info and 1%, 0.1%, and 0.01%
      iii. Put 90ul LB in two microcentrifuge tubes (tubes A and B)
      iv. Move 10ul bacterial mix to 1% plate.
      v. Move 10ul of bacterial mix into tube A. Mix well, then move 10ul from tube A to 0.1% plate.
      vi. Move 10ul from tube A and add to tube B. Mix well, then move 10ul from tube B to 0.01% plate.
      vii. For non-controls, add remaining 990ul of bacterial mix to 50ml LB in flask. Put flask on 37C shaking incubator overnight.
      viii. Use glass beads to spread bacteria on each plate, then put in 37C incubator overnight.
9) The next day, count colonies on all plates to estimate CFUs for full 50ml cultures. Subtract CFUs calculated from negative controls to account for background colonies from undigested backbone plasmid. Estimate fold coverage of variants in the pool. Aim for 10x coverage or better. If lower coverage, repeat electroporation and combine DNA after extraction proportional to coverage, the coverage is additive.
10) Extract DNA from 50ml cultures using Purelink HighPrep Plasmid Midiprep kit (Invitrogen) per kit protocol.
11) Mix pools proportional to how many variants are in each pool to create final pre-barcode plasmid pool.
12) Digest 5ug pre-barcode plasmid pool with 5ul BbsI-HF (New England Biolabs) in 5ul NEB cutsmart buffer and 35ul ddH2O at 37C for 3 hours.
13) Run on 1% agarose gel and extract linearized plasmid Qiaquick Gel Extraction kit (Qiagen) using kit protocol.
14) Purify digest using 0.66x volume Highprep magnetic beads using manufacturer protocol. Elute into 43ul ddH2O.
15) Digest again with BbsI-HF (to digest any remaining uncut backbone).
    a. Mix 43ul hifi product, 2ul BbsI, 5ul NEB cutsmart buffer
    b. Incubate at 37C for 3 hours
16) Purify digest using 0.66x volume Highprep PCR cleanup beads (MagBio) using manufacturer protocol. Elute into 20ul buffer EB.
17) Set up Gibson assembly to add synthesized barcode fragment using NEBuilder HiFi DNA assembly master mix (New England Biolabs) Also set up a negative control reaction combination using ddH2O in place of barcode fragment.:
    a. 100ng linearized plasmid pool (0.014pmol)
    b. 3.3ng barcode fragment (0.05pmol)
    c. 10ul NEBuilder master mix
    d. ddH2O to 20ul total
    e. incubate at 50C for 1 hour
18) Purify Gibson assembly using 0.66x volume Highprep PCR cleanup beads (MagBio) and elute into 20ul buffer EB
19) Electroporate as in step 8.
20) Count colonies as in step 9. Repeat electroporations as needed until total estimate CFUs reaches 115x number of variants in the library (aiming for 115 barcodes/variant means ~95% variants should have 100+ barcodes if normally distributed)
21) Extract DNA from 100ml culture using Purelink HighPrep Plasmid Midiprep kit (Invitrogen) per kit protocol. Prepare final mixture for lentivirus production.

**MPRA Part 4: Plasmid pool barcode association sequencing**

Starting with final MPRA library pool mixture.

1) PCR 1
   a. Plasmid pool	(40ng)
   b. 5’bridge-R1-backbone primer (100uM)	1uL
   c. 3’bridge-GFP (100uM)	1uL
   d. 2x NEB Next PCR MM	100uL
   e. ddH2O	to 200ul total

Split into 5 tubes. Run 98C 60s > [98C 15s, 70.9C 20s, 72C 3mins]x5 > 72C 5mins

2) Purify Highprep PCR cleanup beads (MagBio) using manufacturer protocol. Elute into 50ul buffer EB.
3) PCR 2
   a. PCR 1 product	50uL
   b. P5-iQ50##-5’bridge (100uM)	1uL
   c. P7-iQ70##-NNN-3’bridge (100uM)	1uL
   d. ddh2o	48uL
   e. 2x NEB Next PCR master mix	100uL

Split into 5 tubes. Run 98C 60s > [98C 15s, 70.3C 20s, 72C 3mins]x10 > 72C 5mins.

4) Add 200ul 6x loading dye and run on a 1.8% agarose gel at 100V for an hour.
5) Cut out amplified band and extract DNA using Qiaquick Gel Extraction kit (Qiagen). Elute into an appropriate volume for the amount of gel melted.
6) Purify using Highprep PCR cleanup beads (MagBio) using manufacturer protocol. Elute into 50ul buffer EB.
7) Sequence with high-throughput long-read sequencing.

**MPRA Part 5: Cell Culture**

iPSCs were cultured in mTeSR Plus (Stem Cell technologies) and verified free of mycoplasma using MycoAlertTM (Lonza). Cardiomyocytes were differentated using WNT modulaton as previously described.^1,2^ Cardiomyocyte differentaton was optmized using RPMI plus B27 supplement (Gibco) without insulin during day 0–2 (CHIR 4–6 µM and 0-1 mM actvin A during first 24 h) followed by CDM3 media (without B27) with IWP4 5 µM (Stemgent) during days 3–4. Retnol inhibitor (BMS 453, Cayman Chemical, 1 µM) was included for days 3–6 to minimize atrial lineage differentiation.^3^ IPSC-CMs were maintained in CDM3 until spontaneous contractions. After spontaneous contractions were visible, cardiomyocytes were purified in CDM3-lactate using glucose-deprived, lactate containing media on days 12–16 with RPMI without glutamine (Biologic Industries)^1,2^ and cryopreserved. Differentiation of gene-edited iPSCs were performed within 15 passages of initial gene editing and last karyotyping.

Purified, cryopreserved iPSC-CMs were thawed and replated as monolayers (400,000 cells/cm2) for 8 additional days (approximately day 24 post differentiation) on stiff PDMS membranes (Specialty Manufacturing, Inc.) with growth factor reduced Matrigel (Corning). For MYBPC3 variant RNA and protein experiments, iPSC-CMs were maintained in CDM3-lactate media (4 days) and then oxidative phosphorylation promoting RPMI media (OxPhos) as previously described (Tsan et al.^1^) plus B27 supplement (4 days). Ox-Phos media, adapted from Correia et al.^3^, is based on glucose-free RPMI with 1× B27 supplement and galactose, lactate, glutamax, and pyruvate at final media concentrations of 4, 4, 2, and 0.5 mM, respectively (all from Sigma).

**MPRA Part 6: DNA/RNA extraction from CMs**

1) Collect each well of ~1e6 CMs in 350ul buffer RLT with beta-mercaptoethanol per Allprep DNA/RNA extraction kit (Qiagen). Pool collected cells.
2) Run collected cells through Qiashredder columns (Qiagen) at 10,000+ rpm for 1 minute.
   a. Extract DNA/RNA from shredded cells using Allprep DNA/RNA extraction kit as directed, eluting RNA into 40ul and DNA into 50ul.

**MPRA Part 7: Reverse Transcriptase Reaction**

Using Omniscript reverse transcriptase kit (Qiagen) with targeted primer.

Make multiple 20ul reactions based on volume of extracted RNA.

For each reaction:

RNA	up to 12.75ul

10X Buffer RT	2ul

dNTPs	2ul

RNasin Ribonuclease Inhibitor (Promega)	0.25ul

Primer P7tag-UMI-GFP Primer (10uM)	2ul

Omniscript RT	1ul

ddH2O	to 20ul total (12.75 – RNA)

Incubate at 37C for 1 hour.

**MPRA Part 8: Amplification for Barcode Count Sequencing**

Multi-step amplification to avoid barcode jackpotting. PCRs using NEBNext High Fidelity PCR Master Mix (New England Biolabs)

1) PCR 1
   a. DNA

Extracted DNA	up to 196ul

Primer P5tag-spacer (100uM)	2ul

Primer P7tag -UMI-GFP (100uM)	2ul

ddH2O	196ul-DNA volume

2X NEB High Fidelity PCR MM	200ul

Split into 8 tubes.

b. RNA

Reverse transcriptase product	up to 196ul

Primer P5tag-spacer (100uM)	2ul

Primer P7tag-UMI-GFP (100uM)	2ul

ddH2O	196ul-RNA volume

2X NEB High Fidelity PCR MM	200ul

Split into 8 tubes.

Run both DNA and RNA PCRs: 98C 1min -> [98C 10s -> 69.8C 30s -> 72C 60s]x15 cycles -> 72C 5mins

2) Purify each using Highprep PCR cleanup beads (MagBio) using manufacturer protocol. Elute into 60ul buffer EB.
3) qPCR to determine cycle number for PCR 2: Set up 3 pcr reactions: DNA PCR1 product, RNA PCR1 product, and a negative control using water instead of PCR1 product. Use one of the index primer pairs you plan to use for PCR2.

PCR1 product/ddH2O	5ul

10X SYBR Green	2ul

Primer P7-iQ70##-P7tag (10uM)	1ul

Primer P5-iQ50##-P5tag (10uM)	1ul

ddH2O	1ul

2X NEB High Fidelity PCR MM	10ul

Run 98C 1 min -> [98C 10s -> 67C 30s -> 72C 60s]x30 cycles

Cycle number for PCR2 for DNA and RNA should be selected to be a few cycles before the multicomponent plot curves plateau. Typically, 10-14 cycles.
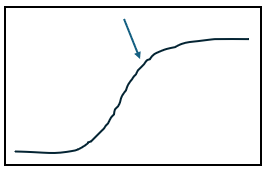

4) PCR 2 – separate reactions for DNA and RNA, may need to run separately if different cycle numbers are needed. Pick index primers to be unique from all others you are planning on sequencing together. Record which primer pairs are used for DNA and which for RNA.

PCR1 product	50ul

Primer P7-iQ70##-P7tag (100uM)	1ul

Primer P5-iQ50##-P5tag (100uM)	1ul

ddH2O	48ul

2X NEB High Fidelity PCR MM	100ul

Split into 5 tubes.

Run 98C 1 min -> [98C 10s -> 67C 30s -> 72C 60s]x?? cycles (based on qPCR)

5) Purify each using Highprep PCR cleanup beads (MagBio) using manufacturer protocol. Elute into 20ul buffer EB.
6) Add 20ul 6x loading dye and run on 1.8% agarose gel at 100V for 1 hour.
7) Cut out band and extract DNA using Qiaquick Gel Extraction kit (Qiagen) using kit protocol. Elute into 50ul buffer EB.
8) Purify each using Highprep PCR cleanup beads (MagBio) using manufacturer protocol. Elute into 20ul buffer EB. THIS IS YOUR FINAL PRODUCT FOR SEQUENCING.
9) Run ~20-30ng of each sample on a 1.8% agarose gel at 100V for 1 hour for final quality check.
10) Mix at 3:1 RNA:DNA molar ratio and submit for high-throughput short-read Illumina sequencing.

**MPRA Barcode Association and Dictionary Creation**

PacBio long-read sequencing reads were processed for barcode dictionary creation on the Great Lakes Computing Cluster. Circular consensus sequencing reads in BAM format were converted to FASTQ using samtools v1.17^4^. Adapter trimming and sequence filtering were then performed using cutadapt v4.4^5^ with a maximum permitted error rate of 10% and a quality trim threshold of q=10 applied to both ends. Variant sequences and barcodes were isolated and partitioned into separate FASTQ files using an Awk script, then matched to the reference library using a custom python script adapted from Cooper et. al. 2022^6^, producing a barcode dictionary file containing variant names and their paired barcodes.

**MPRA DNA/RNA Barcode Counting**

Barcode counting was performed on the Great Lakes Computing Cluster. Raw FASTQ files from Illumina-sequenced DNA and RNA pools were processed using cutadapt v4.4^5^ for adapter trimming and filtering with a maximum permitted error rate of 10% and a quality trim threshold of q=10 applied to both ends. Barcodes were isolated and partitioned into separate FASTQ files using an Awk script. Barcode counting was performed using a custom python script that performed a three-way join among the DNA barcodes, RNA barcodes, and the previously generated barcode dictionary file, discarding barcodes that did not appear in all three. It then created and saved a table of barcodes matched to their variant names and the number of times they appear in the DNA and RNA barcode files.

**MPRA Data Analysis**

DNA and RNA barcode count files for each replicate were processed and analyzed in R v4.4.1.^7^ Each biologic replicate was processed separately as follows: Variants associated with fewer than 10 unique barcodes, and barcodes with fewer than 10 DNA or RNA counts were discarded. Raw counts were normalized to counts per million using the total library read depth. For each barcode, RNA count / DNA count ratio was calculated. Then, for each variant, the interquartile range (IQR) of RNA/DNA ratios was calculated among all associated barcodes. Barcodes with ratios that fell outside of 1.5xIQR above Q3 or below Q1 were discarded as outliers. Following outlier removal, a mean ratio was calculated for each variant. After all replicates were processed, a final dataset was produced using a full outer join, then filtered for variants with data for at least two replicates.

To determine the statistical significance of variant activity relative to wild-type, barcode-level analysis was performed. For each variant, log_2_ transformed RNA/DNA ratios across all associated barcodes was compared to same for wild-type using a Wilcoxon Rank-Sum test with Benjamini-Hochberg correction.

1	Tsan, Y.-C. *et al.* Physiologic biomechanics enhance reproducible contractile development in a stem cell derived cardiac muscle platform. *Nature Communications* **12** (2021 Oct 25). <https://doi.org:10.1038/s41467-021-26496-1>

2	Burridge, P. W. *et al.* Chemically Defined and Small Molecule-Based Generation of Human Cardiomyocytes. *Nature methods* **11** (2014 Jun 15). <https://doi.org:10.1038/nmeth.2999>

3	Correia, C. *et al.* Distinct carbon sources affect structural and functional maturation of cardiomyocytes derived from human pluripotent stem cells. *Scientific Reports* **7** (2017 Aug 17). <https://doi.org:10.1038/s41598-017-08713-4>

4	Li, H. *et al.* The Sequence Alignment/Map format and SAMtools. *Bioinformatics* **25** (2009/08/15). <https://doi.org:10.1093/bioinformatics/btp352>

5	Martin, M. Cutadapt removes adapter sequences from high-throughput sequencing reads. *EMBnet.journal* **17** (2011/05/02). <https://doi.org:10.14806/ej.17.1.200>

6	Cooper, Y. A. *et al.* Functional regulatory variants implicate distinct transcriptional networks in dementia. *Science* **377** (2022-08-19). <https://doi.org:10.1126/science.abi8654>

7	R: A Language and Environment for Statistical Computing (Vienna, Austria, 2024).
